# *Dnmt3a* knockout in excitatory neurons impairs postnatal synapse maturation and is partly compensated by repressive histone modification H3K27me3

**DOI:** 10.1101/2019.12.20.883694

**Authors:** Junhao Li, Antonio Pinto-Duarte, Mark Zander, Chi-Yu Lai, Julia Osteen, Linjing Fang, Chongyuan Luo, Jacinta D. Lucero, Rosa Gomez-Castanon, Joseph R. Nery, Isai Silva-Garcia, Yan Pang, Terrence J. Sejnowski, Susan B. Powell, Joseph R. Ecker, Eran A. Mukamel, M. Margarita Behrens

**Affiliations:** Department of Cognitive Science, University of California San Diego, La Jolla, CA; Computational Neurobiology Laboratory, Salk Institute for Biological Studies, La Jolla, CA; Genomic Analysis Laboratory, Salk Institute for Biological Studies, La Jolla, CA; Waitt Advanced Biophotonics Core, Salk Institute for Biological Studies, La Jolla, CA; Department of Psychiatry, University of California San Diego, La Jolla, CA; Howard Hughes Medical Institute, Salk Institute for Biological Studies, La Jolla, CA; Department of Human Genetics, University of California, Los Angeles, CA

**Keywords:** Epigenetics, DNA methylation, Dnmt3a, H3K27me3, RNA-seq, excitatory synapse, working memory

## Abstract

Two epigenetic pathways of repression, DNA methylation and Polycomb repressive complex 2 (PRC2) mediated gene silencing, regulate neuron development and function, but their respective contributions are unknown. We found that conditional loss of the *de novo* DNA methyltransferase *Dnmt3a* in mouse excitatory neurons altered expression of synapse-related genes, stunted synapse maturation, and impaired working memory and social interest. Loss of *Dnmt3a* abolished postnatal accumulation of CG and non-CG DNA methylation, leaving neurons with an unmethylated, fetal-like epigenomic pattern at −140,000 genomic regions. The PRC2-associated histone modification H3K27me3 increased at many of these sites, partially compensating for the loss of DNA methylation. Our data support a dynamic interaction between two fundamental modes of epigenetic repression during postnatal maturation of excitatory neurons, which together confer robustness on neuronal regulation.

---

Epigenetic modifications of DNA and chromatin-associated histone proteins establish and maintain the unique patterns of gene expression in maturing and adult neurons (Kundakovic and Champagne, 2015). Neuron development requires the reconfiguration of epigenetic modifications, including methylation of genomic cytosine (DNA methylation, or mC) (Guo et al., 2014; Lister et al., 2013a; Stroud et al., 2017) as well as covalent histone modifications associated with active or repressed gene transcription (Fagiolini et al., 2009; Putignano et al., 2007). While mC primarily occurs at CG dinucleotides (mCG) in mammalian tissues, neurons also accumulate a substantial amount of non-CG methylation (mCH) during postnatal brain development in the first 2-3 weeks of life in mice or the first two decades in humans (Lister et al., 2013a). Accumulation of mCH, and the gain of mCG at specific sites, depend on the activity of the *de novo* DNA methyltransferase DNMT3A (Gabel et al., 2015). In mice, the abundance of *Dnmt3a* mRNA and protein peaks during the second postnatal week (Lister et al., 2013a; Stroud et al., 2017), a time of intense synaptogenesis and neuronal maturation. Despite evidence for a unique role of *Dnmt3a* and mCH in epigenetic regulation of developing neurons, the long-term consequences of *Dnmt3a*-mediated methylation on brain function remain largely unknown (Stroud et al., 2017).

One challenge for investigating the developmental role of *Dnmt3a* has been the lack of adequate animal models. Deleting *Dnmt3a* around embryonic day 7.5 driven by the *Nestin* promoter dramatically impaired neuromuscular and cognitive development and led to early death (Nguyen et al., 2007). This early loss of *Dnmt3a* specifically affects the expression of long genes with high levels of gene body mCA (Boxer et al., 2019; Kinde et al., 2016). By contrast, deletion of *Dnmt3a* starting around postnatal day 14 driven by the *Camk2a* promoter caused few behavioral or electrophysiological phenotypes (Feng et al., 2010), with only subtle alterations in learning and memory depending upon the genetic background (Morris et al., 2014). These results suggest that *Dnmt3a* has a critical role during a specific time window between late gestation and early postnatal life. During these developmental stages, regulated gains and losses of DNA methylation throughout the genome establish unique epigenomic signatures of neuronal cell types (Luo et al., 2017; Mo et al., 2015).

To address the role of *Dnmt3a*-dependent epigenetic regulation in the functional maturation of cortical excitatory neurons, we created a mouse line using the *Neurod6* promoter (Schwab et al., 2000) (Nex-Cre) to delete exon 19 of *Dnmt3a (Okano et al., 1999)*. In this conditional knockout (cKO), *Dnmt3a* is functionally ablated in excitatory neurons in the neocortex and hippocampus starting in mid-to-late gestation (embryonic day E13-15; *Dnmt3a* cKO) (Goebbels et al., 2006). In *Dnmt3a* cKO animals, DNA methylation was substantially disrupted in excitatory neurons, leading to altered behavior and synaptic physiology without early life lethality or overt brain morphological alterations. We generated deep DNA methylome, transcriptome and histone modification data in *Dnmt3a* cKO and control pyramidal cells of the frontal cortex, enabling detailed assessment of the molecular basis of neurophysiological and behavioral phenotypes. We found that the Polycomb repressive complex 2 (PRC2) associated chromatin modification, H3K27me3, increases during postnatal development following loss of DNA methylation in cKO neurons. Our data support a dynamic interaction between two fundamental modes of epigenetic repression in developing brain cells, which together confer robustness on neuronal regulation and prevent more extensive consequences for the loss of DNA methylation.

## Results

### *Dnmt3a* conditional knockout in pyramidal neurons during mid-gestation specifically impairs working memory, social interest, and acoustic startle

Previous studies of *Dnmt3a* KO mice yielded results ranging from little cognitive or health effects (Feng et al., 2010; Morris et al., 2014) to profound impairment and lethality (Nguyen et al., 2007). These studies suggest the developmental timing of *Dnmt3a* loss in neurons may determine the extent of subsequent phenotypes. Here, we took a targeted approach by functionally ablating *Dnmt3a* in cortical pyramidal cells starting during mid-gestation (Figure 1A). We took advantage of the developmental onset of *Neurod6* expression between embryonic day E11 and E13, after the onset of *Nestin* expression (Thompson et al., 2014) but well before the major postnatal wave of reprogramming of the neuronal DNA methylome (Figure S1A) (Lister et al., 2013a). We confirmed the deletion of *Dnmt3a* exon 19 in cortical excitatory neurons of cKO animals (Figure 1B and S1B), and the reduction in DNMT3A protein in whole tissue extracts at early postnatal time points (P5 and P13) when *Dnmt3a* mediated accumulation of mCH normally begins in the frontal cortex (Lister et al., 2013b) (Figure S1C). *Dnmt3a* cKO animals survived and bred normally, without overt morphological alterations in the brain (Figure S1D). We found no impairments in gross motor function in an open field test: cKO mice traveled more during the first 5 minutes of testing (Figure S2A) while performing fewer rearings associated with exploratory interest (Figure S2B). Moreover, cKO mice had no signs of increased anxiety-like behavior on three separate behavioral tests (Figure S2C-E), in contrast with the reported anxiogenic effects of *Dnmt3a* knockdown in the mPFC of adult mice (Elliott et al., 2016). The absence of major impairments in overall health, motor function, or anxiety-like behavior established a baseline for investigating the role of *Dnmt3a* in specific cognitive and social behaviors.

**Figure 1.**
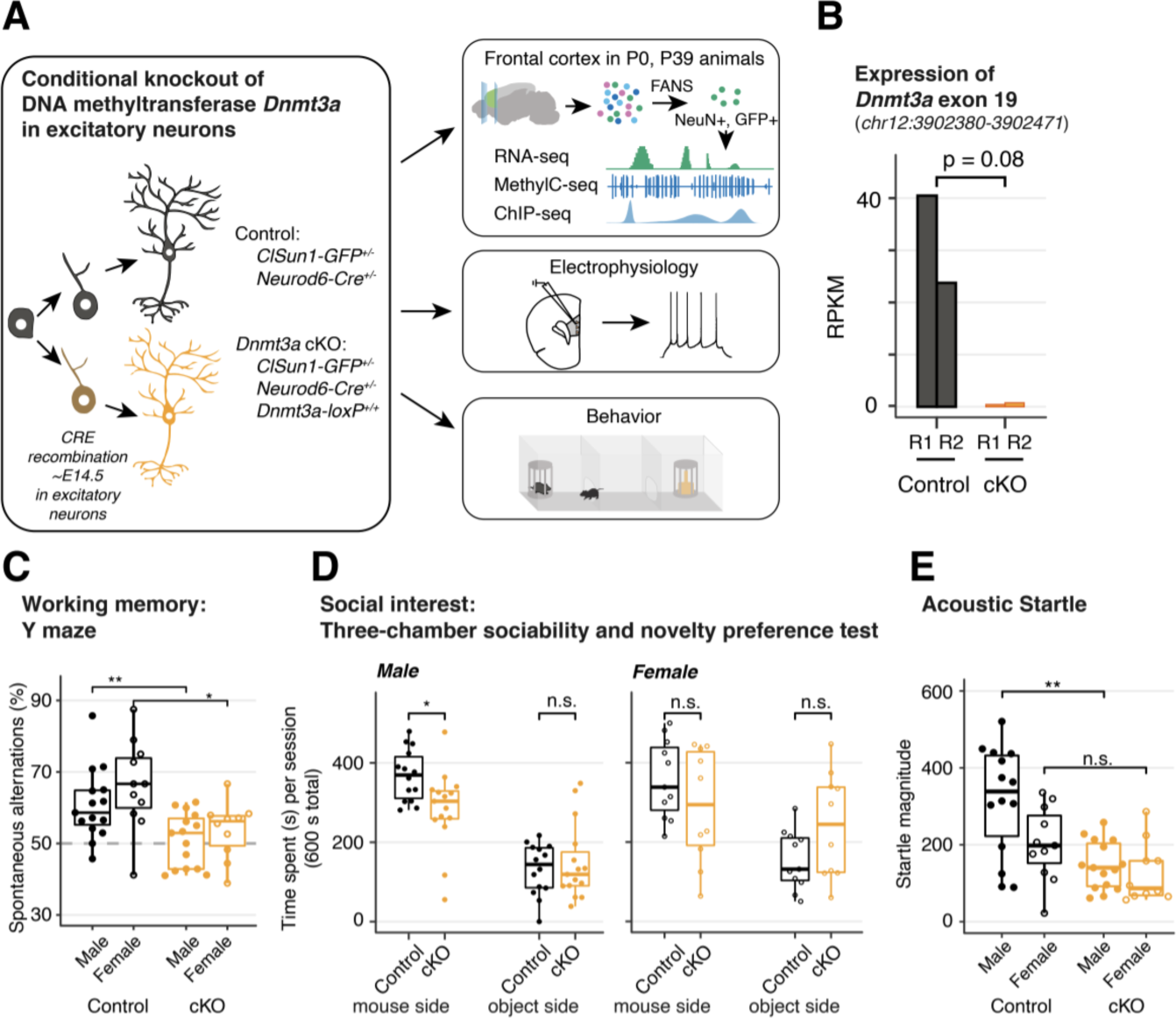
*Dnmt3a* conditional knockout (cKO) in cortical pyramidal neurons during mid-gestation impaired working memory, social interest and acoustic startle responses. **(A)** Experimental model of the conditional loss of *Dnmt3a* in excitatory neurons. P0 and P39, postnatal day 0 and 39. FANS, fluorescence-activated nuclei sorting. **(B)** RNA-seq confirmation of the deletion of *Dnmt3a* exon 19 in P39 excitatory neurons. RPKM, reads per kilobase per million. R1/2, replicate 1/2. T-test p = 0.08. (C) *Dnmt3a* cKO mice made fewer spontaneous alternations in the Y-maze test of working memory (Wilcoxon test, **, p = 0.0079; *, p = 0.011; n =15 male control, 15 male cKO, 11 female control, 10 female cKO). **(D)** Male *Dnmt3a* cKO mice spent less time interacting with an unfamiliar mouse, indicating reduced social interest (Wilcoxon test; * p=0.01048; n =14 male control, 15 male cKO, 11 female control, 10 female cKO) **(E)** Male *Dnmt3a* cKO mice had decreased startle response to a 120 dB acoustic pulse (Wilcoxon test, **, p = 0.0019; n.s., not significant).

We focused on cognitive domains associated with neurodevelopmental illness, including working memory and sensorimotor gating (Habib et al., 2019) and social interest (Dodell-Feder et al., 2015). *Dnmt3a* cKO mice did not alternate spontaneously between the arms of a Y-maze (p = 0.0079 for males, p = 0.011 females, Figure 1C), indicating impaired spatial working memory. Moreover, male *Dnmt3a* cKO animals showed less preference for interacting with a novel mouse in the three-chamber paradigm, indicating reduced social interest (Figure 1D, left panel; p = 0.01048). Male *Dnmt3a* cKO mice also had significantly attenuated acoustic startle reflex (p = 0.0019, Figure 1E and S3B). We observed increased prepulse inhibition (PPI) in male *Dnmt3a* cKO mice, but this may be driven by the reduced startle reflex (Figure S3A, p < 0.05). It is noteworthy that the observed deficits in startle response were not due to hearing deficits, since *Dnmt3a* cKO mice displayed intact hearing in tests of prepulse inhibition and fear conditioning.

To test whether these deficits in specific neuro-cognitive domains reflect generalized impairment in brain function, we assessed long-term memory using a fear conditioning paradigm. There were no significant differences between *Dnmt3a* cKO and control male mice in acquisition or recall of fear memory following re-exposure to the context or conditioned stimulus after 24-48 h (Figure S3C-E), or in extinction (Figure S3F). Altogether, these behavioral results indicate that *Dnmt3a* cKO in excitatory neurons specifically impairs working memory, social interest and acoustic startle.Altered expression of hundreds of genes in *Dnmt3a* cKO excitatory neuronsTo investigate the impact of epigenetic disruption on gene expression, we compared the transcriptomes of cKO and control excitatory neuron nuclei in mature mice (postnatal day 39) (Table S1). We isolated nuclei from excitatory neurons in frontal cortex by backcrossing the *Dnmt3a* cKO animals into the INTACT mouse background (Mo et al., 2015) followed by fluorescence-activated nuclei sorting (FANS) and RNA sequencing. Although sorted nuclei contain only a subset of the cell’s total mRNA and are enriched in immature transcripts, nuclear RNA-seq is nevertheless a quantitatively accurate assay of gene expression that is robust with respect to neural activity-induced transcription (Bakken et al., 2018; Lacar et al., 2016). Nuclear RNA abundance was highly consistent across independent replicates within the same group (Spearman correlation *r* = 0.92-0.95, Figure 2A and S4A). Given the repressive role of DNA methylation (mCG and mCH) in regulating gene expression in neurons (Guo et al., 2014; Lister et al., 2013a), we expected to find increased gene expression in the cKO. Consistent with this, we detected 932 differentially expressed (DE) genes with higher expression in the cKO (FDR < 0.05, Figure 2A and S4B, Table S2). However, we also detected significantly lower expression of 788 genes in cKO neurons, which suggests a complex cellular response to the loss of mC.

**Figure 2.**
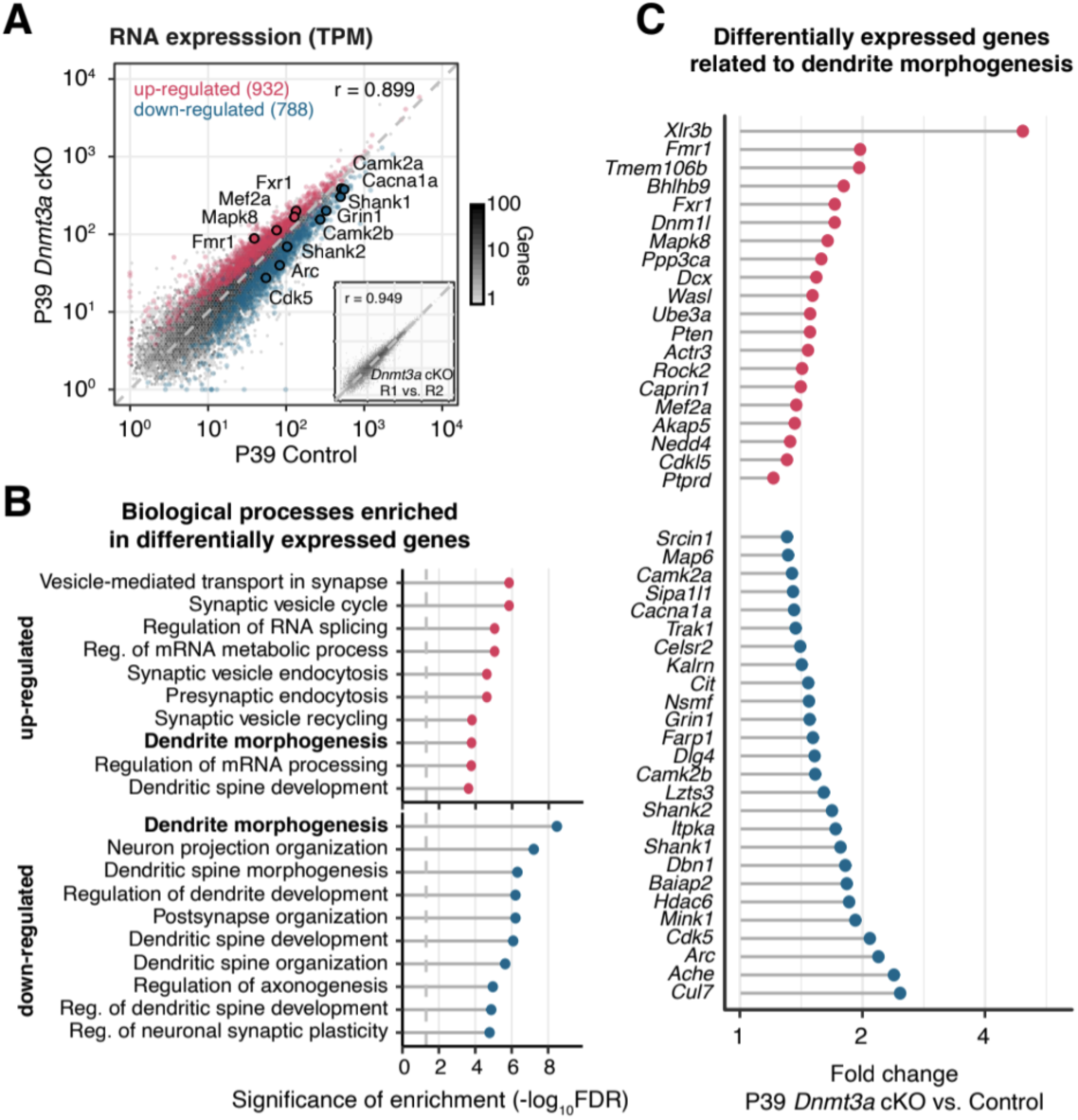
*Dnmt3a* cKO disrupted expression of genes associated with dendritic morphology in cortical pyramidal neurons. **(A)** Gene expression in control vs. *Dnmt3a* cKO excitatory neurons at P39, showing significant up- (red) and down-regulated (blue) differentially expressed (DE) genes (false discovery rate FDR < 0.05). Examples of genes involved in dendrite morphogenesis are highlighted. Inset: Consistent mRNA-seq across two replicates of *Dnmt3a* cKO. TPM, transcripts per million; r, Spearman correlation coefficient. **(B)** Top 10 significantly enriched Gene Ontology Biological Process annotations for DE genes (dashed line: FDR = 0.05). **(C)** Fold-change for DE genes with an annotated role in dendrite morphogenesis.

Many key regulators of synapse function and dendrite structure were differentially expressed in *Dnmt3a* cKO neurons. Both up- and down-regulated genes were significantly enriched in functions related to synaptic physiology and dendrite structure (Figure 2B, FDR<10^-3^, Table S3). Indeed, “dendrite morphogenesis” was the most enriched biological process for down-regulated genes and was also highly enriched among up-regulated genes (Figure 2B and S4C, Table S3). Several DE genes with annotated roles in dendrite morphogenesis were specifically related to glutamatergic synapses and voltage-gated channels, including *Camk2a, Camk2b, Cit, Cacna1a, Shank1* and *Shank2* (Figure 2C). These results suggest that the developmentally regulated mC accumulation in pyramidal neurons contribute to excitatory neuron synapse formation and maturation.Loss of *Dnmt3a* impairs synapse maturation and attenuates neuronal excitabilityTo directly test the impact of *Dnmt3a* cKO on dendritic morphology, we quantified the number and structure of 1,278 DiI-labeled dendritic spines (NexCre/C57: n = 701 from 5 mice; cKO: n = 577 from 4 mice) of layer 2 pyramidal neurons of the mouse prelimbic region (−2 mm anterior to Bregma) (Figure 3A; Methods), a region critical for working memory (Yang et al., 2014) and social approach behavior (Lee et al., 2016). While the overall density of dendritic spines was equivalent in control and *Dnmt3a* cKO neurons (Figure 3B), the spines were significantly longer (mean length 2.219±0.052 µm in cKO, 1.852±0.034 µm in control) and narrower (mean width 0.453±0.008 µm in cKO, 0.519±0.008 µm in control) in *Dnmt3a* cKO neurons (Figure 3C, KS test p < 0.001; Figure S5). Consistent with this, a larger proportion of spines in *Dnmt3a* cKO mice were classified as immature filopodia, and fewer were mushroom-shaped mature spines (Figure 3D) according to pre-established morphometric criteria (see Methods). The proportion of spines with other morphologies, including branched spines with more than one neck (not shown), were not significantly different between genotypes (Figure 3D). These data indicate a role for *Dnmt3a* in dendritic spine maturation.

**Figure 3.**
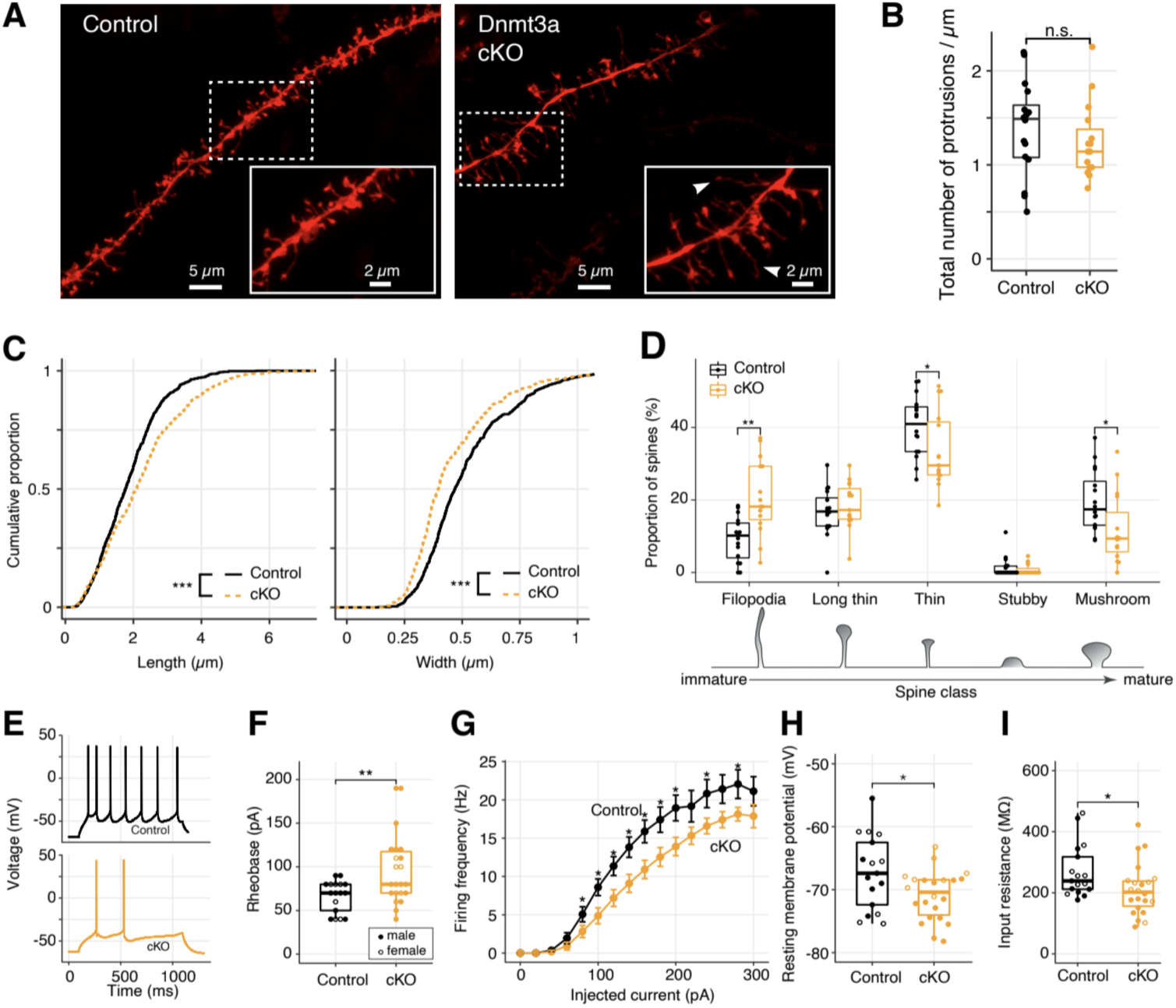
Immature spine morphology and reduced excitability of layer 2 excitatory neurons following *Dnmt3a* cKO. **(A)** Example dendritic segments of layer 2 pyramidal neurons in the prelimbic region labeled with DiI and visualized using a 63x objective coupled to an Airyscan confocal microscope. Arrowheads show filopodia, which were more abundant in *Dnmt3a* cKO mice. **(B)** The density of membrane protrusions was unchanged in the *Dnmt3a* cKO (Wilcoxon test, n.s., not significant). **(C)** Membrane protrusions were significantly longer and narrower in the *Dnmt3a* cKO (K-S test, p < 0.001). **(D)** More spines were classified as immature filopodia, and fewer as mature mushroom shaped spines with large postsynaptic densities (Wilcoxon test, **, p = 0.0015; *, p = 0.046 and 0.011 for Thin and Mushroom, respectively). **(E)** Example whole-cell patch clamp recordings from prelimbic layer 2 pyramidal neurons following 60 pA current injections, the minimal current necessary to trigger an action potential (rheobase) in *Dnmt3a* cKO. **(F)** The median rheobase was significantly higher in the *Dnmt3a* cKO (t-test, **, p = 0.0042). **(G)** Action potential frequency vs. injected current (mean ± s.e.m) showed reduced excitability in *Dnmt3a* cKO (Wilcoxon test, *, p < 0.05). **h,i** *Dnmt3a* cKO neurons were slightly hyperpolarized at V_rest_ when compared to control (Wilcoxon test, *, p = 0.049) and had lower membrane resistance (Wilcoxon test, *, p = 0.023).

We next performed patch-clamp experiments in visually identified layer 2 pyramidal neurons from the prelimbic region to test how the loss of *Dnmt3a* and its impact on spine morphology affected intrinsic neuronal excitability and synapse sensitivity(Figure 3E). Whole-cell current clamp recordings showed that *Dnmt3a* cKO neurons (n = 22 cells from 11 mice) required greater current injections than control (n = 17 cells from 12 mice) to trigger an action potential (higher rheobase, t-test p = 0.0042, Figure 3F), though there was no difference in membrane potential at the firing threshold (Figure S6A). *Dnmt3a* cKO neurons also produced fewer spikes in response to injected current (Figure 3G). These neurons were slightly hyperpolarized at rest (mean −70.90 ± 0.8 mV in cKO, n = 22 cells from 11 mice vs. −67.22 ± 1.4 mV in control, n = 17 cells from 12 mice, Wilcoxon test p = 0.049, Figure 3H), which could reflect differential expression of ion channels at the plasma membrane. Consistent with this, *Dnmt3a* cKO neurons had a lower input resistance (Wilcoxon test p = 0.023, Figure 3I), suggesting increased expression of functional transmembrane ion channels. Whole-cell voltage-clamp recordings of miniature excitatory postsynaptic currents (mEPSCs) showed slight, yet significant, increased amplitude variability in *Dnmt3a* cKO mice (Figure S6B, 8.77±0.32 pA in cKO, n = 8 cells from 4 mice vs. 8.67±0.089 pA in control, n = 8 cells from 5 mice, F-test, p = 0.0032), consistent with a disruption at postsynaptic sites. However, we found no alteration in the mean amplitude (Figure S6B) or frequency (Figure S6C) of mEPSCs recorded at the soma.

### *Dnmt3a* cKO abolishes postnatal DNA methylation

Deletion of *Dnmt3a* during mid-gestation should disrupt the subsequent gain of DNA methylation at specific genomic sites during development (He et al., 2017; Lister et al., 2013a), without affecting sites that maintain or lose methylation after E14.5. Using single base-resolution, whole-genome methylC-seq (Lister et al., 2008), we confirmed that non-CG DNA methylation (mCH) in excitatory neurons is absent at birth (<0.1% of all CH sites at P0), and accumulates by postnatal day 39 (2.11% at P39) (Lister et al., 2013a). The cKO all but eliminated mCH (<0.1% at P0 and P39) (Figure 4A and S7A). While mCH increases in neurons during postnatal life, the genome-wide level of mCG in the brain remains high throughout the lifespan (Lister et al., 2013b). We found that the genome-wide mCG level decreased by 11.8% in mature (P39) cKO neurons (67.8% in cKO vs. 79.6% in control). There was no difference in mCG in newborn mice (P0, 73.1% in both cKO and control, Figure 4A). mCG at P39 was reduced in 61% of all genomic bins (10kb resolution), and was significantly lower in all genomic compartments (promoters, gene bodies, exons, introns, and intergenic regions) (Figure S7B). The reduction in mCG was strongly correlated with reduced mCH (Spearman correlation 0.677, p < 10^-3^, Figure S7C). These data support a role for *Dnmt3a* in postnatal *de novo* CG and CH DNA methylation across genomic compartments (Lister et al., 2013a; Stroud et al., 2017).

**Figure 4.**
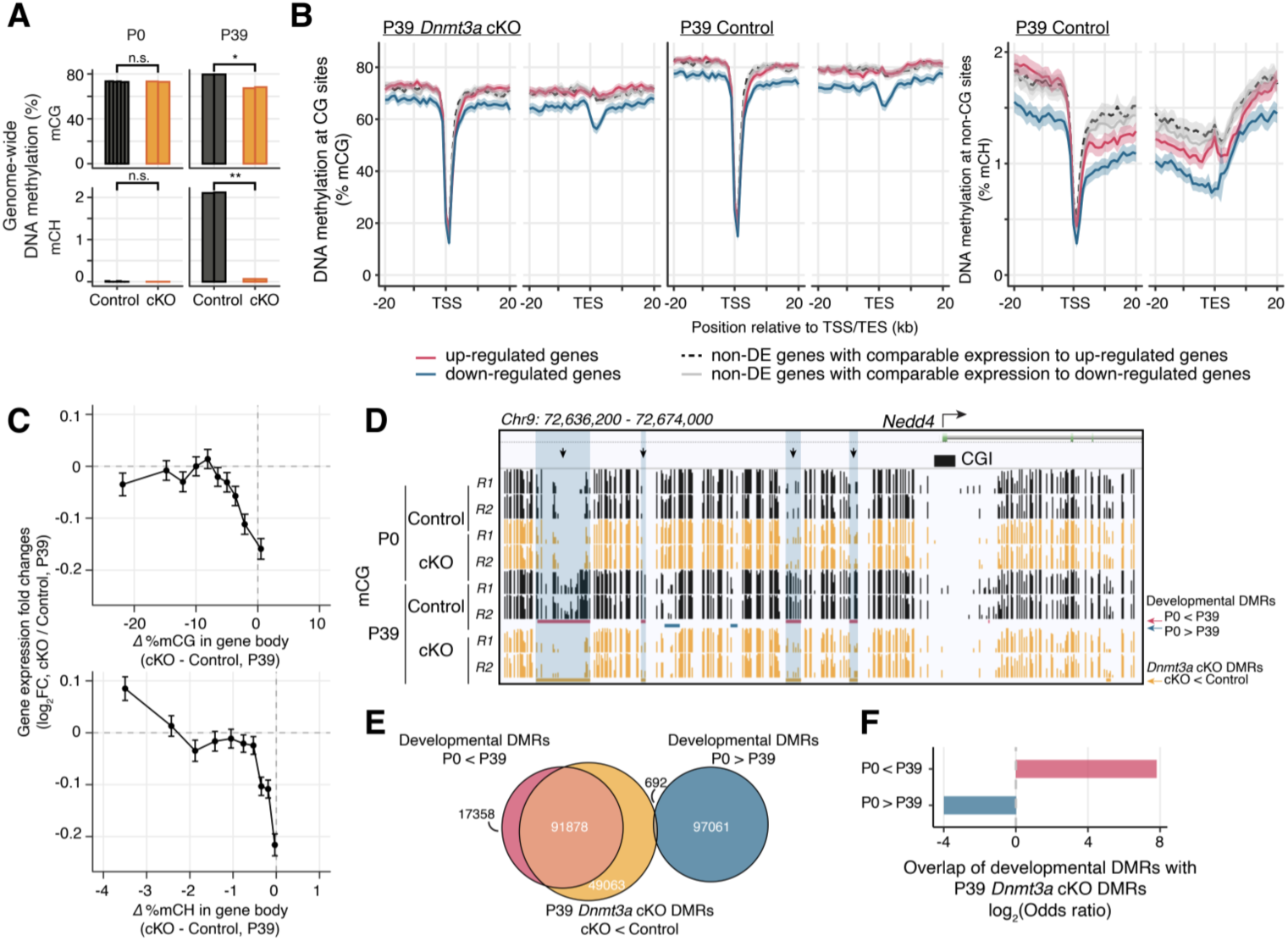
Loss of *Dnmt3a* leaves thousands of genomic regions in a fetal-like demethylated state. **(A)** Non-CG DNA methylation (mCH) is eliminated, and mCG is reduced, in P39 *Dnmt3a* cKO pyramidal cells, while mCG and mCH levels are not changed in P0 (T-test: *, p = 0.025; **, p = 0.0020, n.s., not significant). P0 and P39, postnatal day 0 and 39, respectively. Each bar represents the methylation level in one replicate. **(B)** DNA methylation in P39 pyramidal cells (mCG, mCH; mean ± SEM) in 1 kb bins in the region around the transcription start (TSS) and end site (TES) of DE genes and non-DE genes with matched expression level. **(C)** Difference in gene body methylation vs. fold-change of gene expression between P39 *Dnmt3a* cKO and control. The plots show mean ± SEM gene expression fold-change for genes in 10 non-overlapping bins (deciles of mC difference). **(D)** The *Nedd4* promoter locus contains four differentially methylated regions (DMRs) with naive, fetal-like mCG in P39 *Dnmt3a* cKO. Ticks show mCG at CG sites. Overlapping P39 Dnmt3a cKO DMRs and developmental DMRs are shaded in blue and marked with arrows. CGI, CpG island. R1 and R2, replicates 1 and 2. **(E)** Overlap of P39 *Dnmt3a* cKO DMRs and developmental DMRs. **(F)** P39 *Dnmt3a* cKO hypo-DMRs are significantly enriched (depleted) in DMRs that normally gain (lose) methylation during development (Fisher test, p < 0.05).

### Reduced DNA methylation does not fully explain altered transcription in *Dnmt3a* cKO

We investigated whether the altered gene expression in *Dnmt3a* cKO neurons correlated with loss of DNA methylation at specific sites. We first analyzed DNA methylation around DE genes in mature neurons (P39). The simple model of DNA methylation as a repressive regulator of gene expression predicts that cKO neurons would have lower levels of mC near promoters and gene bodies of up-regulated genes, and higher mC levels at down-regulated genes. Instead, we found the mCG level was lower in cKO neurons around both up- and down-regulated genes (Figure 4B), consistent with the global reduction in mCG in the cKO at P39 (Figure 4A). The difference in gene body methylation (mCG and mCH) was negatively correlated with gene expression changes, consistent with repressive regulation (Gabel et al., 2015; Lavery et al., 2020) (Figure 4C). However, this correlation accounted for <1% of the variance of differential gene expression (Figure S7E).

We next sought to determine if up- and down-regulated genes differ in ways that could explain their different responses to the loss of *Dnmt3a*. Up-regulated genes were on average longer than down-regulated genes (Figure S4D, Wilcoxon, p<10^-5^), consistent with the reported enrichment of mCA and MeCP2-dependent gene repression in long genes (Boxer et al., 2019; Gabel et al., 2015; Kinde et al., 2016). However, there was a broad distribution of gene lengths for both up- and down-regulated genes, and up-regulated genes were not significantly longer than unaffected, non-DE genes. We compared DNA methylation at up- vs. down-regulated genes to test whether they had different epigenetic profiles. In both cKO and control animals, the amount of mCG was similar around up-regulated genes and non-DE genes with comparable expression levels. By contrast, the mCG levels around down-regulated genes were lower than around up-regulated genes and non-DE genes with matched expression (Figure 4B). This suggests that down-regulated genes may be less susceptible to direct repression by DNA methylation, and that their downregulation is an indirect consequence of the loss of mC.Whereas mature *Dnmt3a* cKO neurons lost 11.8% of the normal CG methylation, non-CG methylation was entirely abolished and hence may be more strongly correlated with differential gene expression. Indeed, we found that mCH is more abundant in up-regulated than in down-regulated genes in control neurons (Figure 4B and S7D). However, we also observed lower mCH in both up- and down-regulated genes compared with non-DE genes with equivalent expression levels in control animals. This suggests that some genes with relatively high levels of mCH and mCG in control neurons can maintain their expression despite the loss of DNA methylation, and that de-repression is not a universal outcome for strongly methylated genes. The relatively lower mCH level in down-regulated genes, by contrast, could make them less sensitive to the loss of *Dnmt3a*. The dysregulation of their expression may be due to secondary effects subsequent to the direct loss of DNA methylation.

In addition to promoters and gene bodies, distal regulatory elements such as enhancers are major sites of dynamic DNA methylation where epigenetic regulation can activate or repress expression of genes over long genomic distances through 3D chromatin interactions (Malik et al., 2014). We investigated gene regulatory elements by identifying differentially methylated regions (DMRs) where mCG is altered in cKO compared to control neurons. We found a limited number of DMRs in newborn mice (P0: 1,087 DMRs with lower, 164 with higher mCG in cKO; ≥30% mCG difference; FDR<0.01). In mature neurons (P39), by contrast, we found 141,633 DMRs with substantially lower mCG in cKO compared with controls (Figure S7F; Table S4). Only 19 DMRs had ≥30% higher mCG in cKO. To illustrate, we found five DMRs in a −40 kb region around the promoter of the differentially expressed gene *Nedd4* (Figure 4D). Several of these DMRs were also unmethylated in excitatory neurons in newborn mice, showing that the loss of *Dnmt3a* blocked the developmental gain of mCG at these sites.

The majority of P39 cKO DMRs (67.8%) were distal (≥ 10kb) from annotated transcription start sites. These DMRs were significantly enriched in both active enhancers and in repressed chromatin, suggesting they have a regulatory role (Figure S7G; see also below). We found 109,236 developmental DMRs which lose mCG between P0 and P39 in control neurons (≥30% difference in mCG, FDR<0.01). These DMRs strongly overlapped (84.1%) with the cKO DMRs in mature neurons (Figure 4E-F, Table S4). Moreover, the P39 cKO DMRs were enriched in DNA sequence motifs of multiple transcription factors associated with neuronal differentiation, including *Rest*, *Lhx6*, *Pou3f2* and *Pax6* (FDR<0.05, Figure S7H, Table S5). These results suggest that *Dnmt3a* is essential for the methylation and subsequent repression of neuronal enhancers that are active during prenatal brain development.

Notably, the DMRs in *Dnmt3a* cKO neurons represent only a part of the global reduction in mCG that we observed throughout the genome. Indeed, we found that mCG is reduced by −10% in all genomic compartments, even after excluding P39 DMRs (Figure S7B). Moreover, the density of P39 cKO DMRs around the DE genes was not significantly different in the up- and down-regulated genes and the non-DE genes (Figure S7I). This further supports a complex link between reduced DNA methylation and the transcriptomic changes we observed in the *Dnmt3a* cKO animals.

### Increased PRC2-associated repressive histone modification H3K27me3 in *Dnmt3a* cKO

Given the weak correlation between changes in DNA methylation and gene expression, we explored other potential regulators of the *Dnmt3a* cKO DE genes. To identify transcription factors (TFs) and chromatin regulators with experimental evidence of binding at cis-regulatory regions of the DE genes, we performed Binding Analysis for Regulation of Transcription (BART) (Wang et al., 2018). Chromatin regulators associated with Polycomb repressive complex 2 (PRC2), including *Ezh2*, *Suz12*, *Eed* and *Jarid2*, were among the top DNA binding proteins enriched near the promoters of both up- and down-regulated DE genes (Figure 5A). Several TFs associated with chromatin organization, including the histone deacetylase (*Hdac*) and demethylase (*Kdm*) families, *Cxxc1* and *Ctcf*, were also enriched (Figure S8A, Table S6). These results suggest *Dnmt3a* cKO impacts the chromatin landscape in excitatory neurons, potentially via altered PRC2 activity.

**Figure 5.**
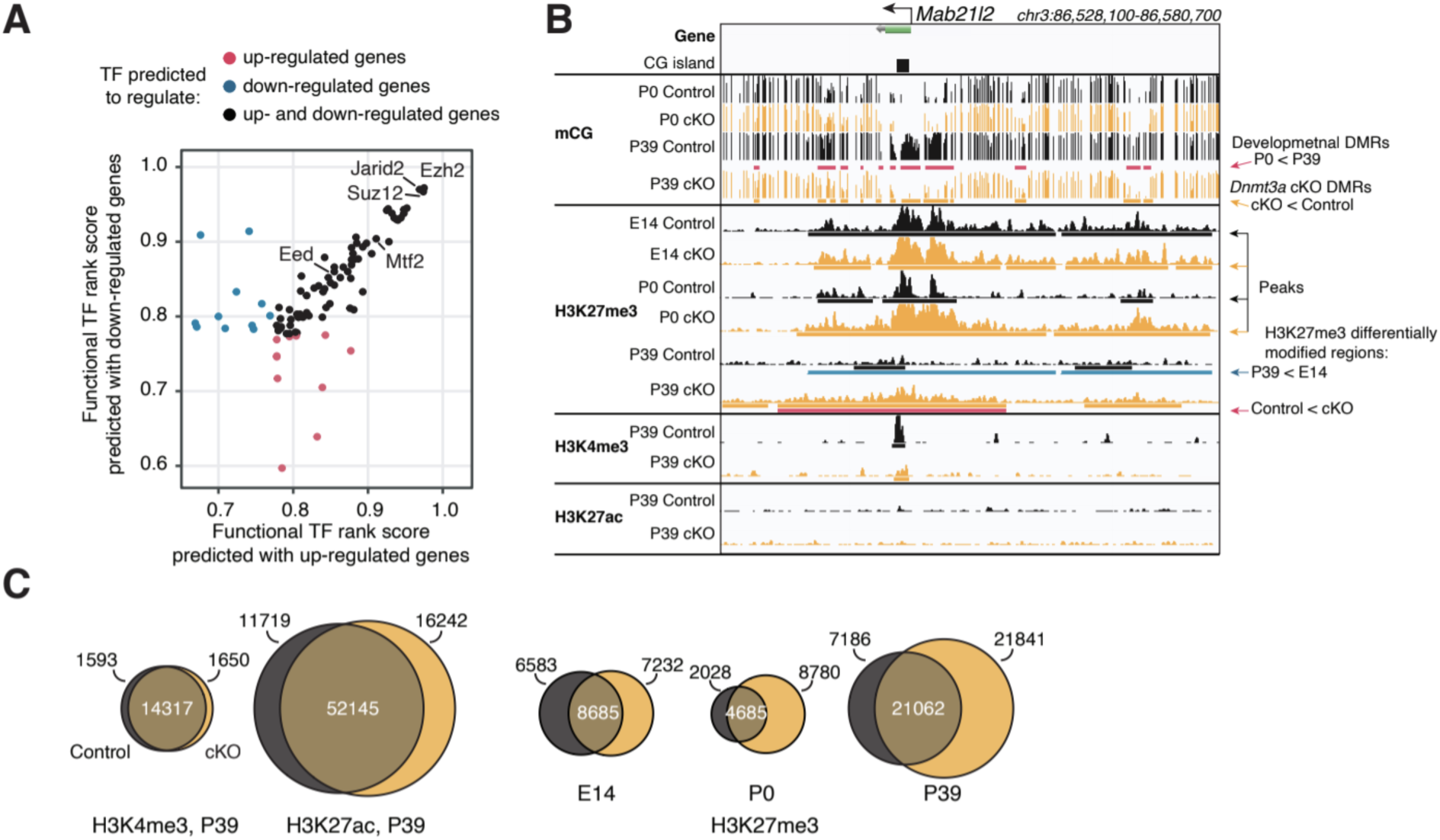
Polycomb repressive complex 2 (PRC2) associated histone modification H3K27me3 is upregulated following loss of DNA methylation. **(A)** Transcription factors (TFs) predicted to regulate P39 *Dnmt3a* cKO differentially expressed genes include many proteins associated with PRC2. The functional TF rank score was assigned by Binding Analysis of Regulation of Transcription (BART)(Wang et al., 2018). (**B)** Browser view of the *Mab21l2* locus, where increased H3K27me3 (differentially modified regions, bottom red bars) coincides with loss of DNA methylation (*Dnmt3a* cKO DMRs, orange bars under the “P39 cKO” track) in P39 *Dnmt3a* cKO. This region loses H3K27me3 during normal development in control pyramidal neurons (blue bars, P39 < E14). DMR, differentially methylated region; E14, embryonic day 14; P0 and P39, postnatal day 0 and 39. **(C)** Histone modification ChIP-seq peaks for active marks (H3K4me3, H3K27ac) are largely preserved in the *Dnmt3a* cKO, while repressive H3K27me3 peaks expand.

To experimentally address this, we performed ChIP-seq in excitatory neurons at embryonic day 14 (E14) and postnatal days 0 and 39 to measure trimethylation of histone H3 lysine 27 (H3K27me3), a repressive mark whose deposition is catalyzed by PRC2 that is important for transcriptional silencing of developmental genes. In P39 neurons, we also measured two histone modifications associated with active chromatin: H3K4me3 (trimethylation of histone H3 lysine 4, associated with promoters) and H3K27ac (acetylation of histone H3 lysine 27, associated with active promoters and enhancers) (Heinz et al., 2015). The active and repressive marks, as well as mCG and mCH, had positive and negative correlations with mRNA expression, respectively (Figure S8B). We noted regions, such as the *Mab21l2* locus (Figure 5B), where increased H3K27me3 in P39 *Dnmt3a* cKO neurons coincided with the loss of methylation at DMRs.

Using a conservative strategy to call ChIP-seq peaks (Zang et al., 2009), we found that marks associated with transcriptional activity (H3K4me3 and H3K27ac) were largely conserved in the P39 cKO and control (Figure 5C). By contrast, we found 51.9% more H3K27me3 peaks in mature (P39) cKO than control neurons (Figure 5C, Table S7). When we directly identified differentially modified (DM) regions, we found no DM for H3K4me3 and H3K27ac between the cKO and control in P39 neurons (Figure S8C). By contrast, we found 4,040 regions with significantly increased H3K27me3 in the P39 cKO, covering −31.05 MB of the genome (Figure S8D, FDR<0.05, Table S8). Differential H3K27me3 appears late during brain maturation: only 3 DM regions were found in earlier development stages (E14 or P0; Figure S9). Genes associated with these DM regions were enriched in development related functions (Figure S8E, Table S9). These results suggest that the increase of H3K27me3 in *Dnmt3a* cKO excitatory neurons occurred post-natally, following the major impact of loss of *Dnmt3a* on neuronal DNA methylation.

The increase in H3K27me3 was closely associated with regions that lost DNA methylation in the *Dnmt3a* cKO. The majority (56.1%) of the regions marked by H3K27me3 in the cKO overlap with P39 cKO DMRs (Figure 6A, Fisher test p < 1e-100). Likewise, P39 cKO DMRs were significantly enriched in peaks and DM regions of H3K27me3 (28.1% overlapped H3K27me3 peaks in cKO, p<1e-100) (Figure 6B). The DMRs were less associated with H3K27ac (13.5%, p < 1e-100), and depleted in regions marked by H3K4me3 (1.1%, p = 4.5e-38) (Figure 6A-B and S8F-G). Moreover, H3K27me3 was more abundant at the center of DMRs in cKO compared to control neurons at P39 (Figure 6C). There was no corresponding increase of H3K27me3 at these DMRs in newborn (P0) or fetal (E14) neurons. Going beyond overlaps of regions, we found a quantitative association between the changes in DNA methylation and H3K27me3 in mature (P39) neurons (Figure 6D). At H3K27me3 peaks, the ChIP-Seq signal intensity fold-change between cKO and control correlated with the loss of mCG in cKO (Spearman r = −0.27, p < 0.001).

**Figure 6.**
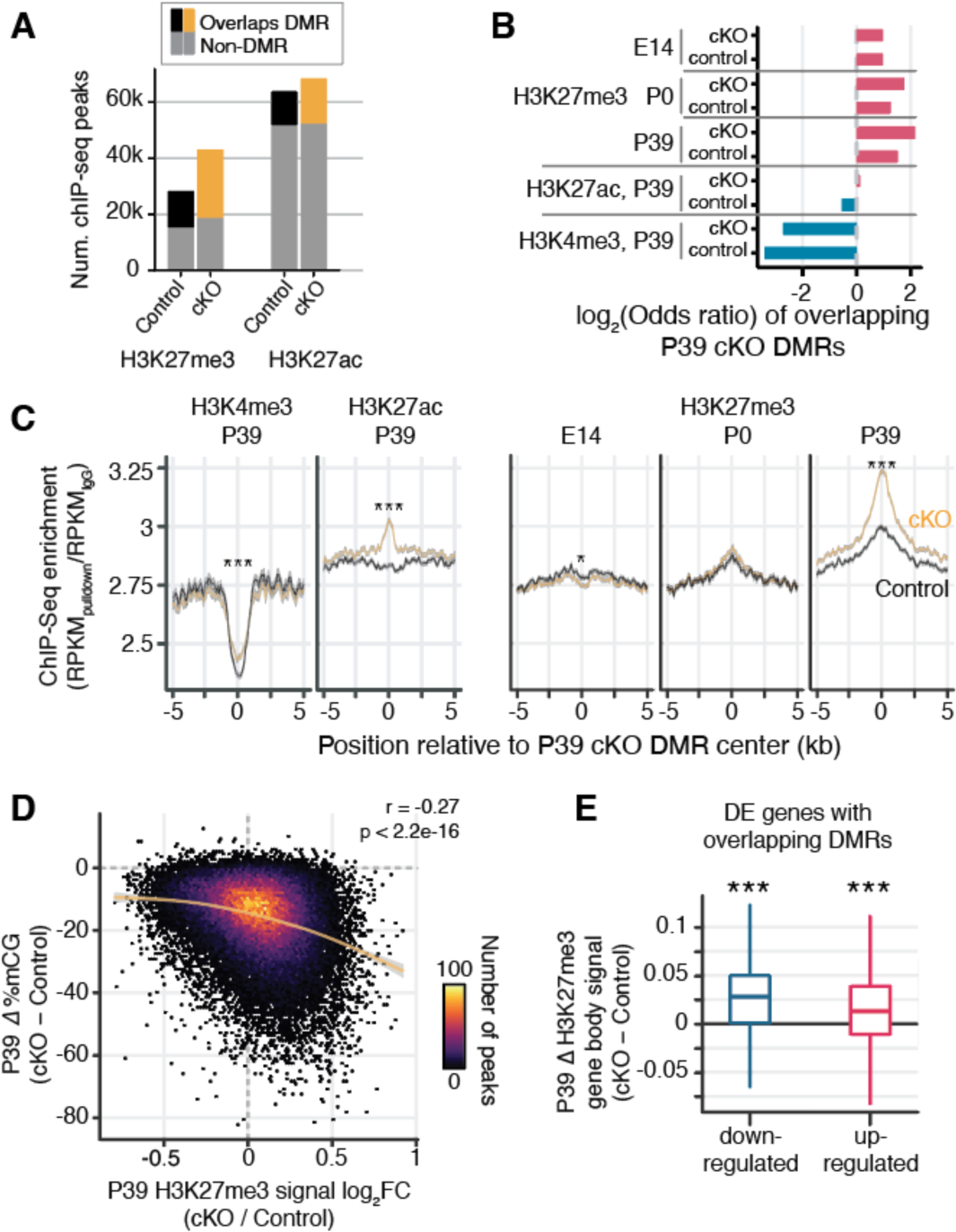
Increased H3K27me3 compensates loss of postnatal DNA methylation. **(A)** Most (56.1%) of the P39 H3K27me3 peaks, but only some (23.3%) of the P39 H3K27ac peaks, overlap with P39 *Dnmt3a* cKO DMRs. **(B)** Significant enrichment (red) or depletion (blue) of P39 *Dnmt3a* cKO DMRs in the histone modification ChIP-seq peaks (Fisher test, p < 0.05). E14, embryonic day 14; P0 and P39, postnatal day 0 and 39. **(C)** Histone modification ChIP-seq signal around the center of DMRs. Shaded ribbon indicates standard error of the mean. RPKM, reads per kilobase per million. ***, Wilcoxon rank sum test p < 0.001. **(D)** Correlation of P39 H3K27me3 signal fold-changes and P39 CG methylation levels differences between *Dnmt3a* cKO and control in H3K27me ChIP-seq peaks. The smoothed line is fitted using a generalized additive model, and the shaded area shows 95% confidence interval of the fit. r, Spearman correlation coefficient. **(E)** P39 *Dnmt3a* cKO DE genes with overlapping P39 *Dnmt3a* cKO DMRs show small but significant increases of H3K27me3 in P39 cKO. ***, Wilcoxon rank sum test against zero, p < 0.001. DE genes that do not overlap with P39 cKO DMRs are shown in Figure S10A. H3K27me3 signal was calculated as the RPKM fold-change between H3K27me3 and IgG.

We further examined whether the increased H3K27me3 could account for the reduced expression of some genes in the *Dnmt3a* cKO neurons. We found that down-regulated genes that overlap DMRs gained 114% more H3K27me3 than up-regulated genes (Fig. 6E, Wilcoxon rank sum test p = 7.35e-6; Figure S10A). These data may partially explain the fact that 788 genes were down-regulated even after the loss of repressive DNA methylation in the *Dnmt3a* cKO (Figure 2A). PRC2-mediated repression may compensate for the loss of mCG and/or mCH, acting as an alternative repressive mechanism when DNA methylation is disrupted.Developmental changes in H3K27me3 are not affected by *Dnmt3a* cKOOur finding that *Dnmt3a* cKO disrupts the normal developmental gain of DNA methylation prompted us to ask whether the changes in H3K27me3 in the cKO are likewise associated with developmental regulation of H3K27me3. Indeed, our ChIP-Seq data from E14, P0 and P39 excitatory neurons revealed striking developmental dynamics in H3K27me3. We identified 12,994 developmentally regulated H3K27me3 DM regions between E14 and P39, with a similar number of regions that gain (6,774) and lose (6,220) H3K27me3 (Figure 7A and S10B, Table S10). Genes associated with these DM regions were enriched in biological processes involved in development such as nervous system development and neurogenesis (Figure S10C). The developmental H3K27me3 DM regions overlapped developmental DMRs, with a notable overlap of regions gaining both mCG and H3K27me3 (Figure 7B). Both modifications may thus act together to repress thousands of genomic regions during development.

**Figure 7.**
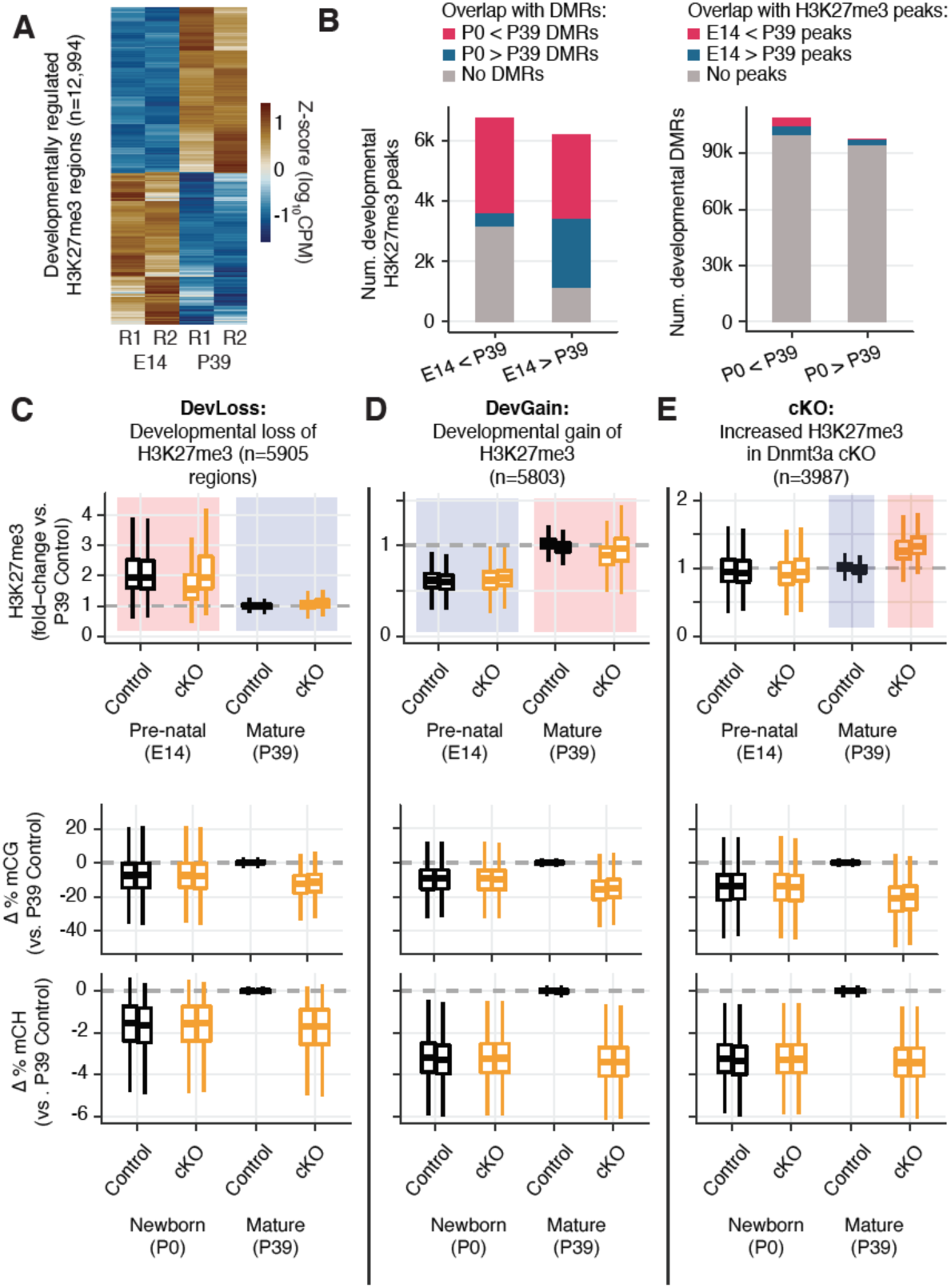
Developmental dynamics of H3K27me3. **(A)** Heatmap of developmentally regulated H3K27me3 regions in E14 and P39 control samples. CPM - counts per million; R1/2 - replicates. **(B)** Barplots show the numbers of developmental differentially modified H3K27me3 regions that overlap developmental DMRs (left panel), and the numbers of developmental DMRs that overlap developmental differentially modified H3K27me3 regions (right panel). **(C-E)** Normalized H3K27me3 signal (fold-changes compared to P39 Control), and mCG, mCH differences (compared to the average of the two replicates from P39 Control) in peaks that overlaps with developmental loss-of-H3K27me3 regions (**C**), developmental gain-of-H3K27me3 regions (**D**), or increased H3K27me3 in P39 *Dnmt3a* cKO (**E**).

The P39 *Dnmt3a* cKO changes of H3K27me3 signal were weakly correlated with the developmental changes of H3K27me3 signal (r = −0.12, p<2.2e-16, Figure S10D). Moreover, there was little appreciable difference between the H3K27me3 signal fold-changes between P39 and E14 cKO samples compared with those of control samples (Spearman r = 0.67, p < 0.001, Figure S10E).

To further stratify the joint distribution of developmental and *Dnmt3a* cKO-dependent changes in both H3K27me3 and DNA methylation, we assigned peaks to three groups (Figure 7C-E and S10F-H). Group DevLoss and DevGain peaks lose or gain H3K27me3 during development, respectively (Figure 7C-D and S10F-G). Group cKO peaks have higher H3K27me3 in the *Dnmt3a* cKO compared to control at P39 (Figure 7E and S10H). We found that developmental peaks (DevLoss and DevGain) were relatively unaffected by the cKO (Δ H3K27me3 = 0.11, −0.20 respectively, in units of log10(CPM+1)), whereas Group cKO had ten-fold larger mean effect (Δ H3K27me3 = 1.12 respectively). Group cKO peaks alsoexperienced greater loss of mCG (Δ mCG = −21.7%) than Group DevLoss (−13.0%) or DevGain(16.3%) (Figure 7C-E and S10F-H, middle and right panels). These results suggest that regions prone to alteration of H3K27me3 by *Dnmt3a* cKO are distinct from the regions affected by developmentally dynamic H3K27me3.

### Novel DNA methylation valleys with increased H3K27me3 signal in the *Dnmt3a* cKO

DNA methylation and H2K27me3 have complementary roles at DNA methylation valleys (DMVs), i.e. large regions (≥5 kb) with low mCG (:515%) that occur around key transcriptional regulators of development in human and mouse tissues (Mo et al., 2015; Xie et al., 2013). Previous studies comparing the epigenetic profile of DMVs across tissues identified multiple categories, including constitutive DMVs present in all tissues as well as tissue-specific DMVs (Li et al., 2018). We found more than twice as many DMVs in P39 cKO (2,969) compared with control (1,351) neurons (Figure 8A, Table S12), covering a greater genomic territory (25.29 MB in cKO, 11.93 MB in control). Most DMVs had active histone marks (H3K4me3+, H3K27me3-), while some had repressed or bivalent profiles consistent with PRC2-associated gene silencing (H3K27me3+) (Figure 8B). By clustering the DMVs using their pattern of DNA methylation, chromatin modifications, and gene expression, we found six distinct categories (Figure 8C-E). Whereas most DMVs lack H3K27me3 (clusters C1, 2, 4 and 5), we found two clusters associated with moderate (C3) or high (C6) levels of H3K27me3. Cluster C6 DMVs, such as one of the promoters of *Foxp2* (Figure 8C), had high H3K27me3 and low mCG in both control and cKO neurons, and were not strongly affected by loss of mC in the *Dnmt3a* cKO. By contrast, cluster C3 DMVs, including the second promoter of *Foxp2* and the promoter of *Slc17a6* (encoding vesicular glutamate transporter, Vglut2), gain mC during normal development (Figure 8C). Some of these DMVs gained H3K27me3 in the P39 cKO compared to control animals (Figure 8E). The loss of mC in these regions in the *Dnmt3a* cKO did not lead to strong activation of gene expression, potentially due to compensatory PRC2-mediated repression.

**Figure 8.**
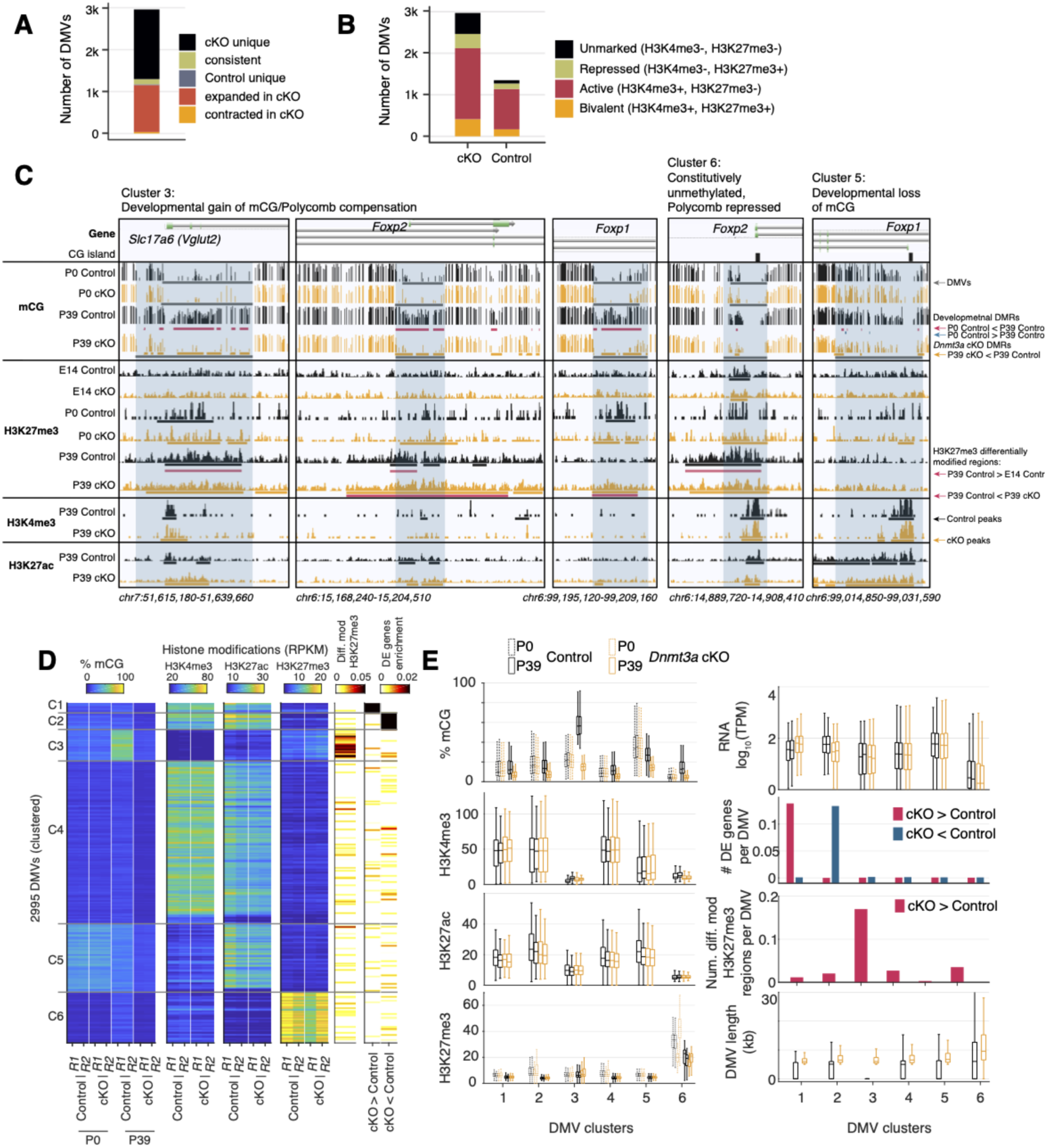
Distinct clusters of DNA methylation valleys were associated with the increased H3K27me3 signal in the *Dnmt3a* cKO. **(A)** Number of DMVs identified in the P39 *Dnmt3a* cKO and the Control samples, categorized by whether they appear in one or both groups or change size in the cKO. **(B)** Overlap of DMVs with the H3K4me4 and/or the H3K27me3 ChIP-seq peaks. **(C)** Browser tracks showing examples of unique DNA methylation valleys (DMVs) in the *Dnmt3a* cKO samples and the increased H3K27me3 signal in their flanking regions. **(D)** Heatmap of DMVs clustered by their methylation levels and histone modifications. The last two columns show the enrichments of DM peaks of H3K27me3 and DE genes. R1/2, replicate 1/2; RPKM, reads per kilobase per million. **(E)** Distribution of DNA methylation, histone marks, gene expression, DE gene enrichment, enrichment of DM peaks of H3K27me3, and the length of the DMVs.

The remaining clusters lack H3K27me3 and are instead marked by low mCG and either high H3K27ac (cluster C5) or both H3K27ac and H3K4me3 (clusters C1, C2). Cluster C5 DMVs, such as the *Foxp1* promoter (Figure 8C), have high mCG in the fetal brain and lose methylation in excitatory neurons during brain development. This demethylation is not affected in the *Dnmt3a* cKO, and we found little difference between the control and cKO neurons at these sites. Finally, clusters C1 and C2 were enriched for up- and down-regulated genes, respectively. Like C5 DMVs, they lose methylation during excitatory neuron development and are expanded in the cKO compared with control animals at 6 weeks of age.

## Discussion

To determine the role of DNA methylation in neurons after their birth, we developed a mouse model where loss of DNA methylation occurs in postmitotic neurons, prior to the postnatal increase in *Dnmt3a* expression and non-CG methylation (Lister et al., 2013a). Conditional deletion of *Dnmt3a* in cortical excitatory neurons abolished non-CG methylation and reduced CG methylation throughout the genome. We found that *Dnmt3a* cKO neurons had both increased and decreased expression of hundreds of genes. Similar complex patterns in gene expression were recently reported when *Dnmt3a* was deleted in inhibitory neurons (Lavery et al., 2020) and following manipulation of the DNA methylation reader MeCP2 (Boxer et al., 2019; Johnson et al., 2017). These gene expression changes may partly reflect the direct effect of lower gene body CH methylation and of loss of CG and CH methylation at gene promoters and distal enhancers (Boxer et al., 2019; Clemens et al., 2019). In addition, some gene expression changes could result from the disruption of other regulatory processes, such as transcription factor expression or chromatin modification. Indeed, we found that the *Dnmt3a* cKO DMRs overlapped with regions that gain methylation during normal post-natal development. These DMRs had increased H3K27ac in the *Dnmt3a* cKO, consistent with observations in adult animals lacking *Dnmt3a* specifically in GABAergic *Sst-* or *Vip-*expressing interneurons (Stroud et al., 2020). In those experiments, embryonic gene-regulatory elements had lower cytosine methylation and increased H3K27ac and H3K4me1 (Stroud et al., 2020). This suggests an essential role for *Dnmt3a* and DNA methylation in shaping the transcriptome during development in part via inactivation of embryonic enhancers, with potentially long lasting effects for the gene expression pattern of mature neurons.

There is a strong antagonistic relationship between DNA methylation and thePRC2-associated histone mark, H3K27me3 (Brinkman et al., 2012; Jermann et al., 2014; Lynch et al., 2012; Reddington et al., 2013; Wu et al., 2010). Switching between Polycomb- and DNA methylation-mediated repression has been observed during development and in cancer (Mohn et al., 2008; Schlesinger et al., 2007; Widschwendter et al., 2007). Severe depletion of mCG can lead to redistribution of H3K27me3, causing derepression of developmental regulators such as the *Hox* gene clusters (Reddington et al., 2013). We did not observe ectopic expression of these genes, possibly due to the relatively modest reduction in mCG in our model compared with cells lacking *Dnmt1.* Instead, we found that in cortical excitatory neurons, thousands of sites gained H3K27me3 following the loss of mCG in the *Dnmt3a* cKO. These sites, which normally gain methylation during postnatal development, were left unmethylated in cKO neurons. These regions were largely distinct from the sites that gain or lose H3K27me3 during normal development. A subset of these regions form large-scale DNA methylation valleys (DMVs) spanning key regulatory genes. Overall, our results suggest that when DNA methylation is disrupted, PRC2-mediated repression may partially compensate for the loss of mCG and/or mCH and act as an alternative mode of epigenetic repression.

We found no differential expression in adult *Dnmt3a* cKO animals of the four core components of PRC2 (*Ezh2*, *Suz12*, *Eed* and *Rbbp4*). However, the expression of *Mtf2* (also known as *Pcl2*), whose protein product is a Polycomb-like protein reported to recruit PRC2 to its target genes (Perino et al., 2018), was −47% higher in the *Dnmt3a* cKO (FDR = 0.016, Supplementary Table 2). *Mtf2* was also one of the top predicted regulators of DE genes based on comparison with public ChIP-Seq datasets (Wang et al., 2018). These findings could indicate a potential mechanism for the increased H3K27me3 in *Dnmt3a* cKO neurons.

Mid-gestation deletion of *Dnmt3a* in pyramidal neurons caused the differential expression of 1,720 genes in excitatory neurons of the frontal cortex of animals analyzed at 5 weeks of age. A substantial fraction of those genes had annotated roles in dendrite morphology and synapse function. Previous studies reported gross motor deficits and a shortened lifespan following early embryonic deletion of *Dnmt3a (Nguyen et al., 2007)*. By contrast, only slight alterations occurred when the deletion occurred past the second postnatal week (Feng et al., 2010; Morris et al., 2014). In contrast, our mid-gestation *Dnmt3a* cKO mice had no overt motor deficits, but were impaired in working memory, social interest and acoustic startle. We showed that layer 2 pyramidal neurons in the mPFC of *Dnmt3a* cKO animals had a larger number of immature dendritic spines and were significantly less sensitive to somatic injections of depolarizing current compared to control neurons. They had lower input resistance, suggesting increased expression of functional transmembrane ion channels. They were also hyperpolarized at rest compared to control, but fired around similar membrane voltages. It is thus noteworthy that *Dnmt3a* cKO mice expressed significantly more mRNA encoding the potassium channel subunit *Kcnq3* (39.5% increase, FDR=0.006, Table S2), which mediates hyperpolarizing M-currents in the sub-threshold voltage range (Brown and Yu, 2000) and regulates resting membrane potential and neuronal excitability (Schwake et al., 2000).

Our study used conditional ablation of *Dnmt3a* in excitatory neurons of the mPFC driven by the NeuroD6 promoter, which is also expressed in the hippocampal dentate gyrus and sub-cortical regions (Goebbels et al., 2006). These regions may contribute to part of the behavioral deficits we observed. However, the specific impairments in working memory and social interest, and in synaptic maturation in layer 2 neurons of mPFC, together suggest that disruption of methylation patterns established by *Dnmt3a* during mPFC neural maturation could have mechanistic relevance in the context of neurodevelopmental disorders. These data complement recent genetic risk association studies linking *Dnmt3a* with autism (C Yuen et al., 2017; Sanders et al., 2015).

Our findings highlight the critical and interconnected roles in brain development and cognitive function of two major modes of epigenetic repression of gene expression: DNA methylation and PRC2-mediated repression. The loss of DNA methylation in excitatory neurons has far-reaching effects on gene expression, synaptic function, and cognitive behavior, and triggers a compensatory gain of the PRC2-associated repressive mark H3K27me3. Although H3K27me3 partially compensates for the loss of DNA methylation, the developmental programs are nevertheless disrupted and lead to altered circuit formation and behavioral dysfunction. Our cKO is a restricted manipulation of one neuron type, yet it directly impacts DNA methylation throughout the genome at millions of sites. Moverover, additional non-enzymatic functions of *Dnmt3a*, other than its methyltransferase activity, may also contribute to the phenotypes we observed. Future work focusing on earlier developmental stages, and using targeted methods to manipulate epigenetic marks in local genomic regions (Liu et al., 2016), may help elucidate the causal interactions among epigenetic modifications that are critical for neuronal maturation and function.

## METHODS

Methods and associated references are available in the online version of the paper.

## Acknowledgments

This work was supported by R01MH112763 to MMB and JRE, and a Kavli Foundation award to MMB, APD and SBP. JRE is an Investigator of the Howard Hughes Medical Institute. We acknowledge stimulating discussions with Huda Zoghbi and Laura Lavery. We thank the members of the Salk Biophotonics Core Dr. Uri Manor, Sammy Weiser Novak, and Dr. Tong Zhang for insightful suggestions. We also thank Joseph Chambers and Colleen Heller for technical assistance in animal handling, and Ms. Faith Zhang for her involvement in the morphometric analysis of dendritic spines. The Waitt Advanced Biophotonics Core Facility at the Salk Institute receives funding from NIH-NCI CCSG: P30 014195 and the Waitt Foundation. The Flow Cytometry Core Facility of the Salk Institute receives funding from NIH-NCI CCSG: P30 014195. The authors have no conflict of interest in relation to the work described here.

## Data access

All sequencing data are available in the Gene Expression Omnibus under accession GSE141587 (reviewer access password: qvuxqsuqrxupxwv). A genome browser displaying the sequencing data is available at https://brainome.ucsd.edu/annoj_private/mm_dnmt3a_ko/ (for reviewer access, username: reviewer, password: dnmt3a).

## Author contributions

JL, APD, JRE, EAM, MMB designed research. MMB, YP, JDL created transgenic mouse models and prepared samples for sequencing. APD, SBP performed and interpreted behavioral studies. APD performed slice electrophysiology studies. APD, JO, CYL, LF, ISG analyzed spine morphology. CL, JRN, RGC, performed methylC-seq, RNA-seq. MZ, JRN and RGC performed ChIP-seq. JL, CL performed bioinformatic and computational analysis of sequencing data. JL, APD, MZ, CL, JRE, EAM, MMB interpreted the findings. JL, APD, EAM, MMB wrote and edited the manuscript in consultation with all authors. MMB, JRE, EAM supervised the research.

---

**Supplementary Figure 1.**
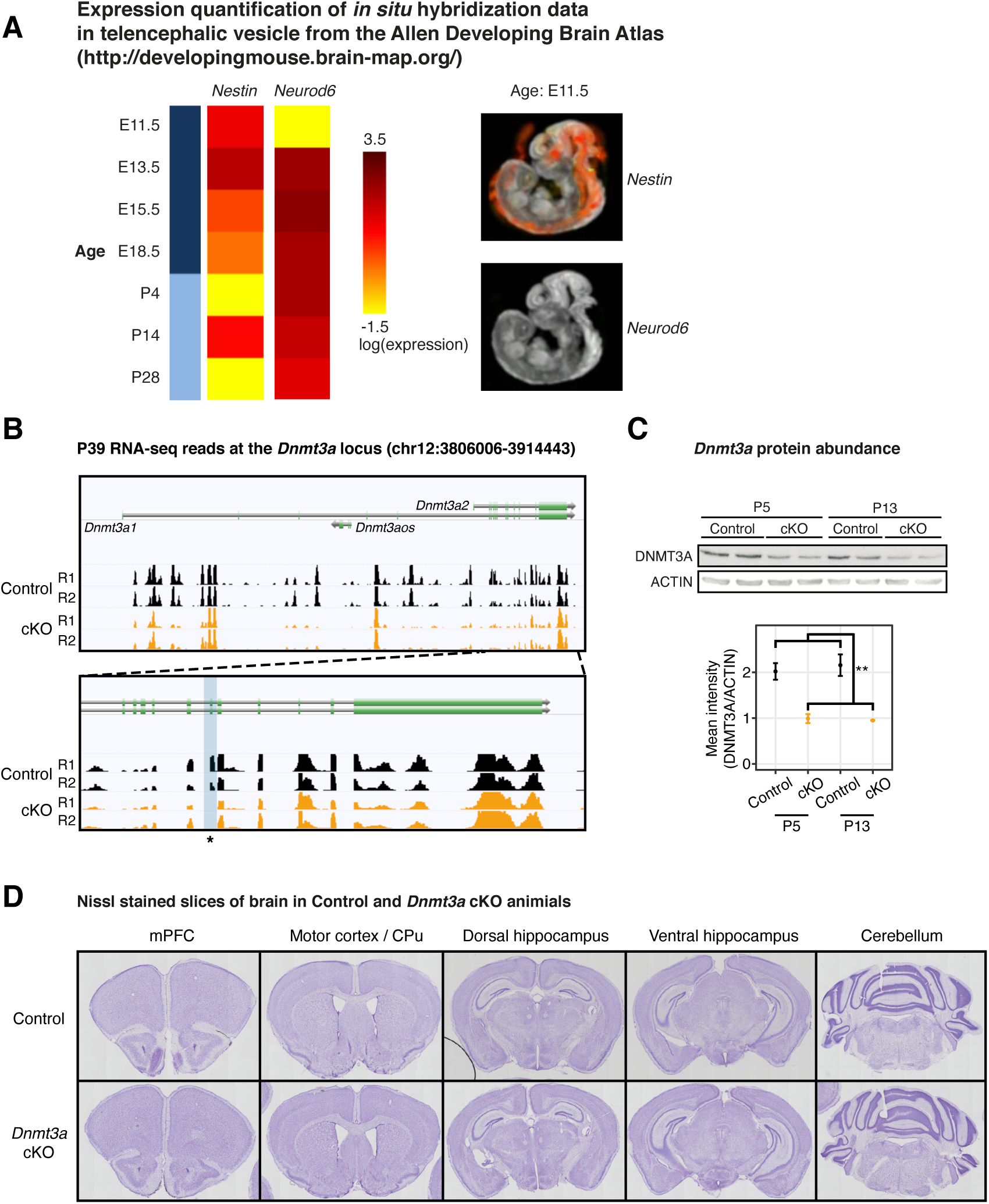
*Dnmt3a* was disrupted on both the mRNA and protein levels in the *Dnmt3a* cKO excitatory neurons. **(A)** Expression quantification of *in situ* hybridization data of gene *Nestin* and *Neurod6* in telencephalic vesicle from the Allen Developing Brain Atlas (http://developingmouse.brain-map.org/). Left panel, heatmap of the gene expression across ages during development. Right panel, example images of the gene expression in E11.5. Image credit: Allen Institute. E11.5-18.5, embryonic days; P4-28, postnatal days. **(B)** Genome browser tracks of mRNA-seq data show confirmation of the deletion of *Dnmt3a* exon 19 in P39 *Dnmt3a* cKO excitatory neurons. The targeted exon region is highlighted in the light blue shaded box with an asterisk. R1/2, replicate 1/2. **(C)** The protein product of the *Dnmt3a* gene is disrupted in the cKO sample. Top panel, Western blot; Bottom panel, quantification of the protein abundance. P5 and P13, postnatal days 5 and 13. **, T-test p = 0.0017. **(D)** Nissl stained slices show no morphological alterations in the brain of the *Dnmt3a* cKO animals. mPFC, medial prefrontal cortex; CPu, caudate putamen.

**Supplementary Figure 2.**
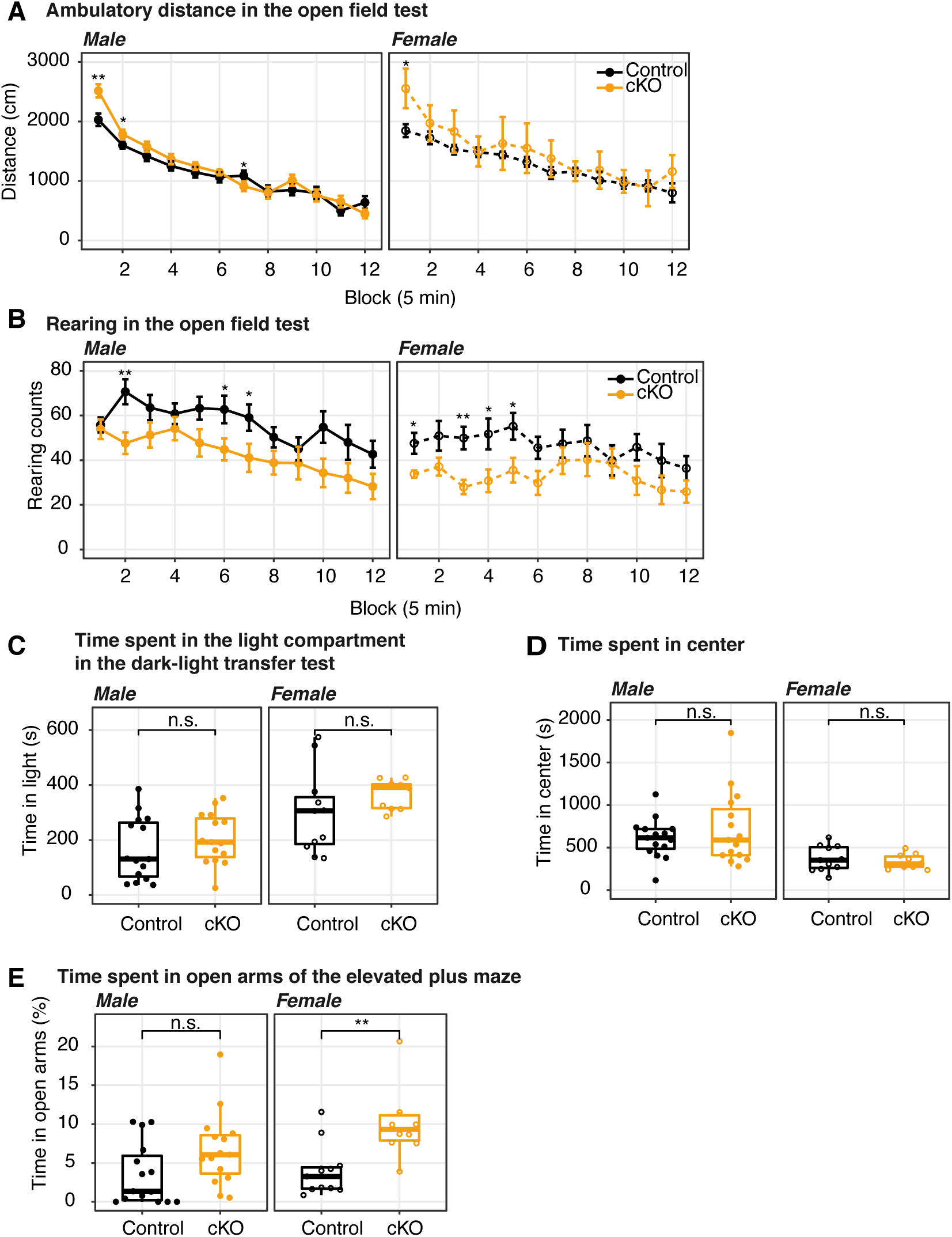
The conditional ablation of *Dnmt3a* in pyramidal neurons did not significantly impair motor activity nor increased anxiety levels. **(A)** and **(D)**, *Dnmt3a cKO* mice displayed normal behavior in the open field test, traveling a similar distance **(A)** as control mice, and also showing a similar degree of center activity **(D)**. **(B)** The exploratory activity was slightly decreased in *Dnmt3a* cKO animals, as suggested by an attenuated rearing behavior. **(C)** The time spent in light in the dark-light transfer test was not significantly affected by the lack of *Dnmt3a.* **(E)** The female, but not the male, cohort of *Dnmt3a* cKO mice spent significantly more time than control mice in the open arms of the elevated plus maze, consistent with lower anxiety levels (Wilcoxon test, **, p = 0.0048; n.s., not significant.). In the line plots **(A-B)**, data were presented as mean ± s.e.m. In all boxplots **(C-E)**, the middle horizontal bar represents the median; the lower and upper hinges correspond to the first and third quartiles, and the whisker extends from the hinge to the value no further than 1.5 * IQR from the hinge, where IQR is the interquartile range. The values of individual experiments are represented by dots superimposed on the boxplots. Wilcoxon test significance: *, p < 0.05; **, p < 0.01; n.s., not significant.

**Supplementary Figure 3.**
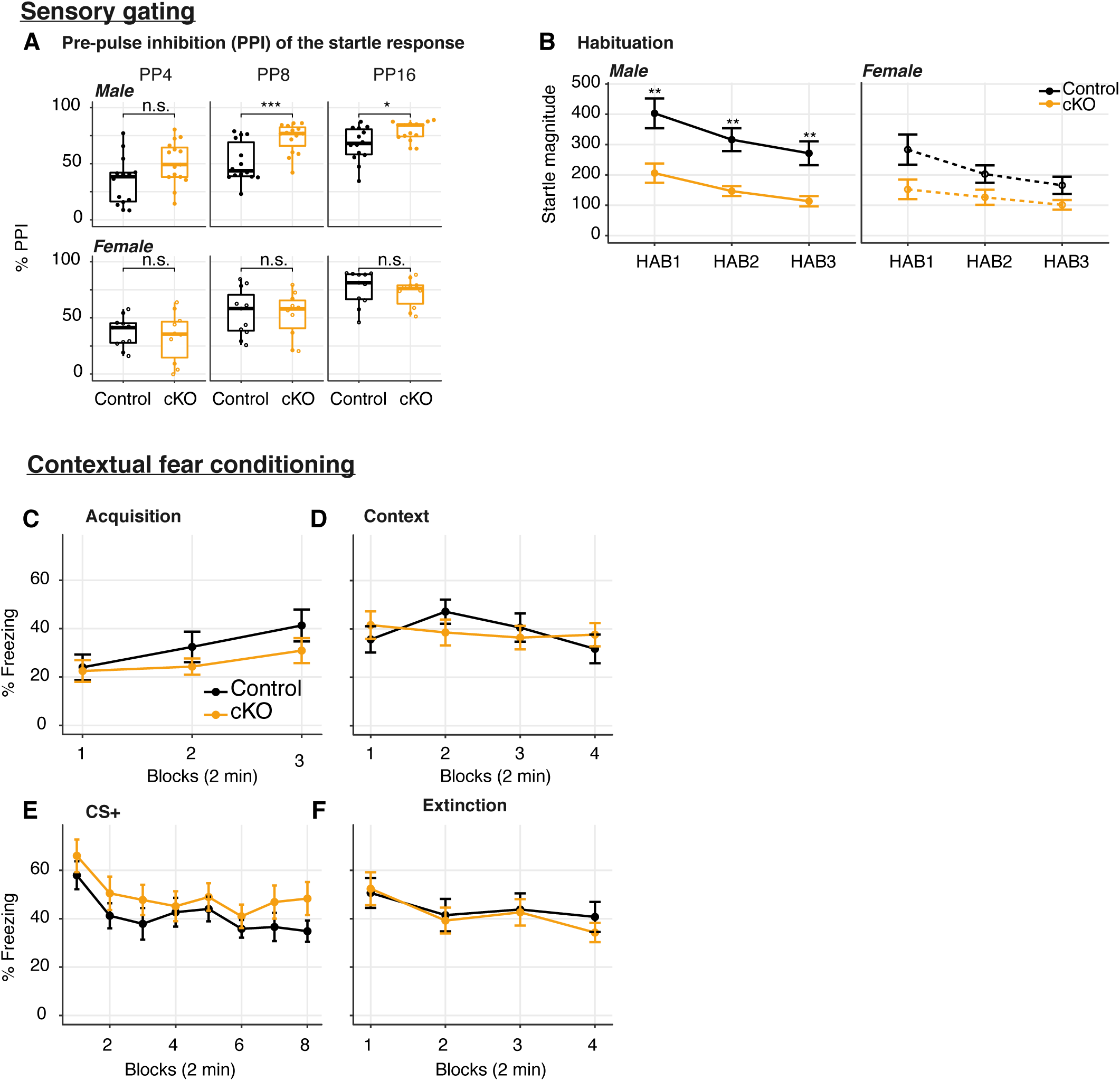
*Dnmt3a* cKO-induced impairment of startle response was accompanied by increased prepulse inhibition, and the cKO did not affect fear memory. **(A)** The percentage of PPI at prepulse intensity of 69, 73 and 81 dB (4, 8 and 16 dB above the 65 dB background, respectively) was increased in male, but not female mice (Wilconxon test, ***, p = 0.00076; *, p = 0.016; n.s, not significant). **(B)** The increased PPI accompanied the impairment in startle responses to a 120 dB tone played at three time points during the recording session (HAB1 - beginning of session; HAB2 - middle of session; HAB3 - end of session) (Wilconxon test p = 0.0027, 0.0019 and 0.0035 in male HAB1, HAB2, HAB3, respectively, and not significant in female). The habituation to the 120 dB auditory tone (i.e. the relative reduction in startle response throughout the experiment) was not significantly different between genotypes. **(C-F)** Fear learning and extinction were tested over four consecutive days. *N* = 14-15 per group. **(C)** Fear acquisition to three tone-shock pairings occurred on day 1; **(D)** contextual fear in relation to the acquisition context (8 min, Block = 2 min) was measured on day 2; **(E)** cued fear recall and extinction training occurred on day 3 (Block = four tone trials), and **(F)** extinction recall (Block = four tone trials) occurred on day 4. Wilcoxon test reported no significant changes between the *Dnmt3a* cKO and control. In boxplots (**A**), the middle horizontal bar represents the median; the lower and upper hinges correspond to the first and third quartiles, and the whisker extends from the hinge to the value no further than 1.5 * IQR from the hinge, where IQR is the interquartile range. The values of individual experiments are represented by dots superimposed on the boxplots. In the line plots (**B-F**), data were presented as mean ± s.e.m.

**Supplementary Figure 4.**
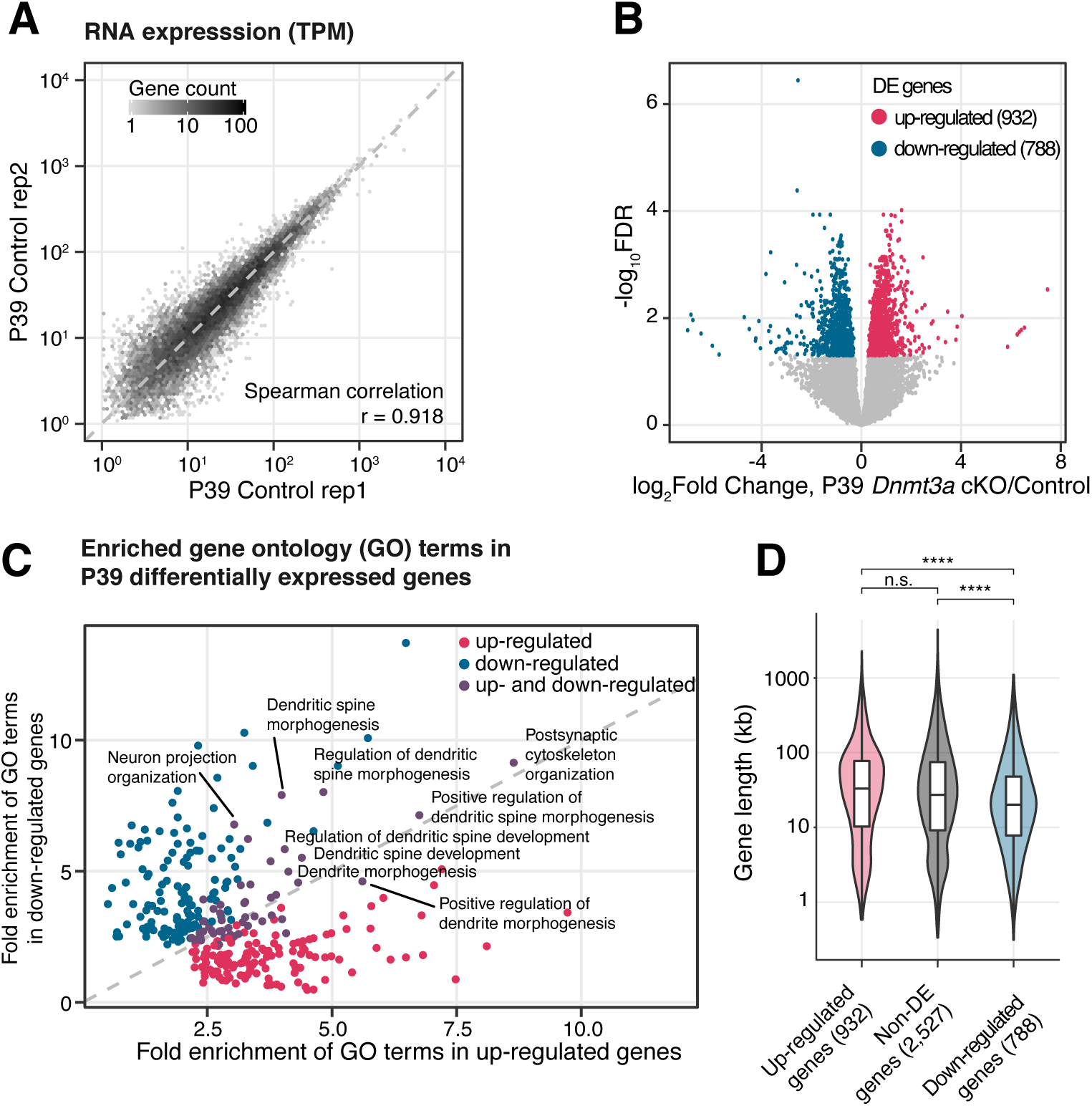
RNA-seq data showed transcriptomic disruption, especially in genes associated with dendritic morphogenesis, in *Dnmt3a* cKO pyramidal neurons. **(A)** Correlation of gene expression in the two biological replicates of the control mouse excitatory neurons. TPM, transcripts per million. r, Spearman correlation coefficient. **(B)** Volcano plot shows the gene expression fold-change of P39 *Dnmt3a* cKO vs. Control samples and their significance. Significant up- and down-regulated differentially expressed genes (DE genes, FDR < 0.05) are colored in red and blue, respectively. **(C)** Comparison of fold enrichment of GO terms significantly enriched (FDR < 0.05) in up- and down-regulated genes in P39. Terms significantly enriched only in up-regulated genes, enriched only in down-regulated genes and enriched in both the up- and down-regulated genes are colored in red, blue and purple, respectively. Representative terms related to dendrite and spine development are labeled. **(D)** Gene length distribution of P39 DE genes. As comparison, non-DE genes were selected with FDR < 0.05 and fold-change < 1.1 (see Supplementary Table 2). The down-regulated genes are generally shorter than the up-regulated genes or the non-DE genes. kb, kilobases. Wilcoxon test, ****, p < 10^-5^, n.s. not significant.

**Supplementary Figure 5.**
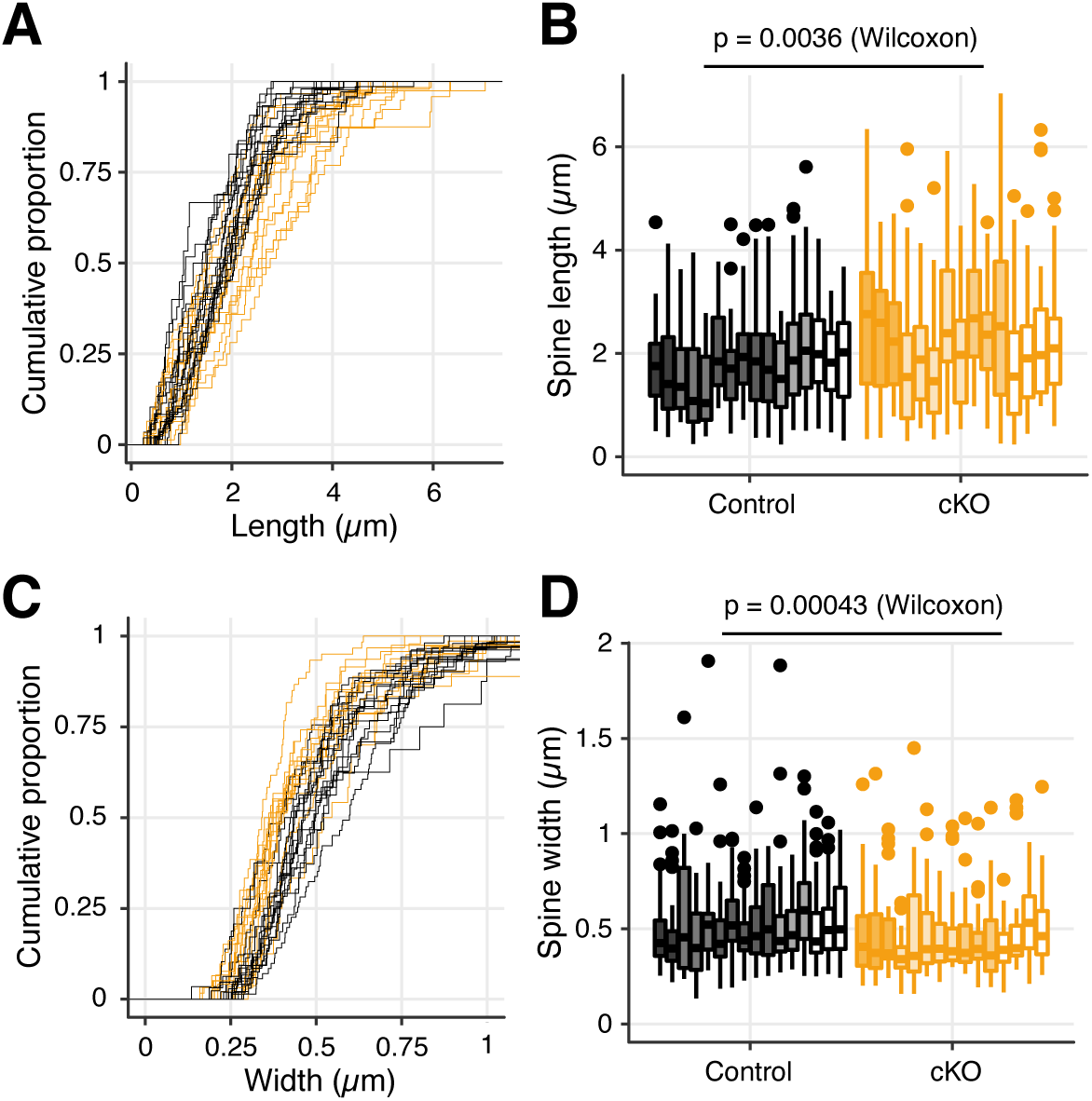
The membrane protrusions in the *Dnmt3a* cKO showed longer dendritic spines and narrower heads. Related to Figure 3C. Here each line/box represents data from one dendrite fragment. **(A)** and **(C),** The cumulative distribution of the length (**A**) and width (**C**) of membrane protrusions in the *Dnmt3a* cKO (orange), as compared to control mice (black). **(B)** and **(D)**, distribution of the length (**B**) and width (**D**) of the protrusions. The shaded colors in the box represent the individual animals from which the dendrite fragment originated (5 control mice and 4 cKO mice). The middle horizontal bar represents the median; the lower and upper hinges correspond to the first and third quartiles, and the whisker extends from the hinge to the value no further than 1.5 * IQR from the hinge, where IQR is the interquartile range. The Wilcoxon test was done with the medians of the boxes (16 control vs. 15 cKO).

**Supplementary Figure 6.**
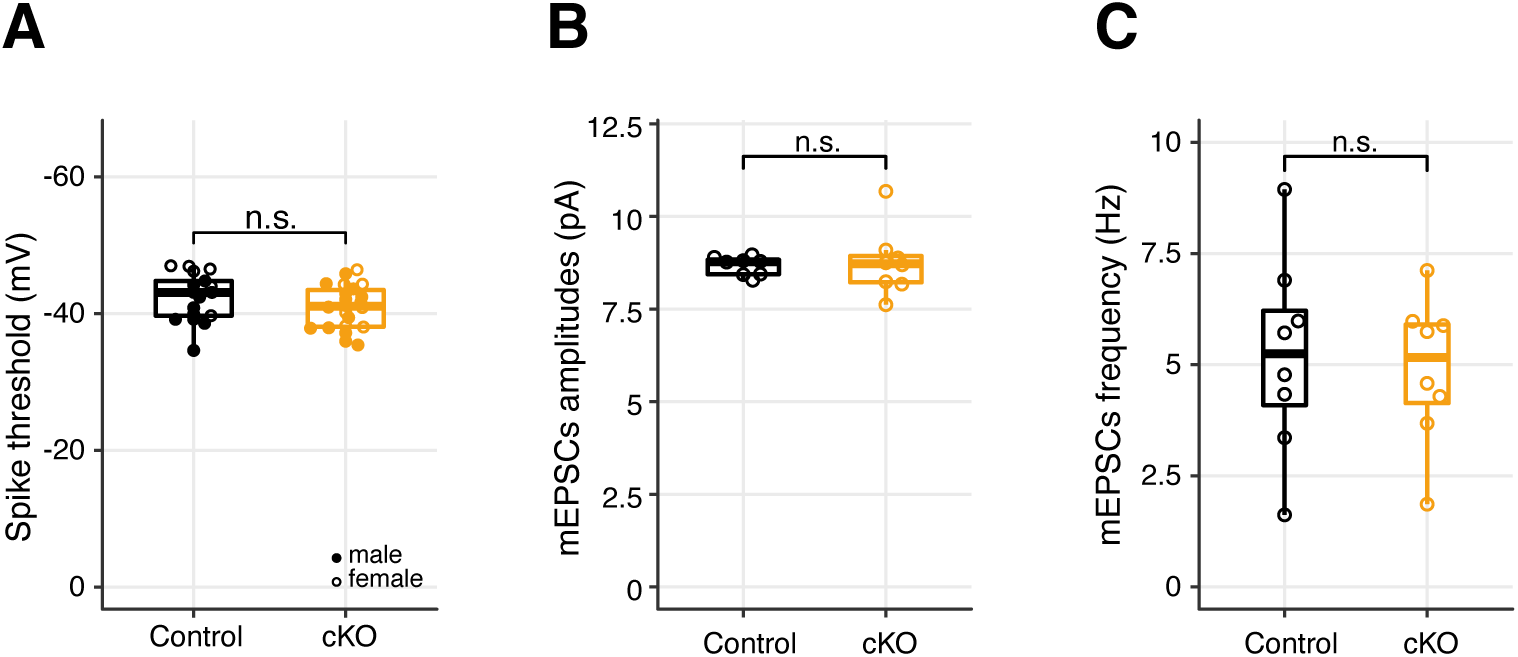
The conditional ablation of *Dnmt3a* in pyramidal neurons did not significantly alter the membrane potential threshold for action potential generation, the mean amplitude or frequency of miniature excitatory postsynaptic events. **(A)** Action potentials were initiated at a similar membrane potential in both genotypes (Wilcoxon test, n.s., not significant). **(B)** While mEPSCs amplitude was not significantly different between genotypes (Wilcoxon test, n.s., not significant), it was slightly, yet significantly, more variable in the *Dnmt3a* cKO (F-test p = 0.0032). **(C)** mEPSCs frequency was not significantly changed. In all boxplots, the middle horizontal bar represents the median; the lower and upper hinges correspond to the first and third quartiles, and the whisker extends from the hinge to the value no further than 1.5 * IQR from the hinge, where IQR is the interquartile range. The values of individual experiments are represented by dots superimposed on the boxplots.

**Supplementary Figure 7.**
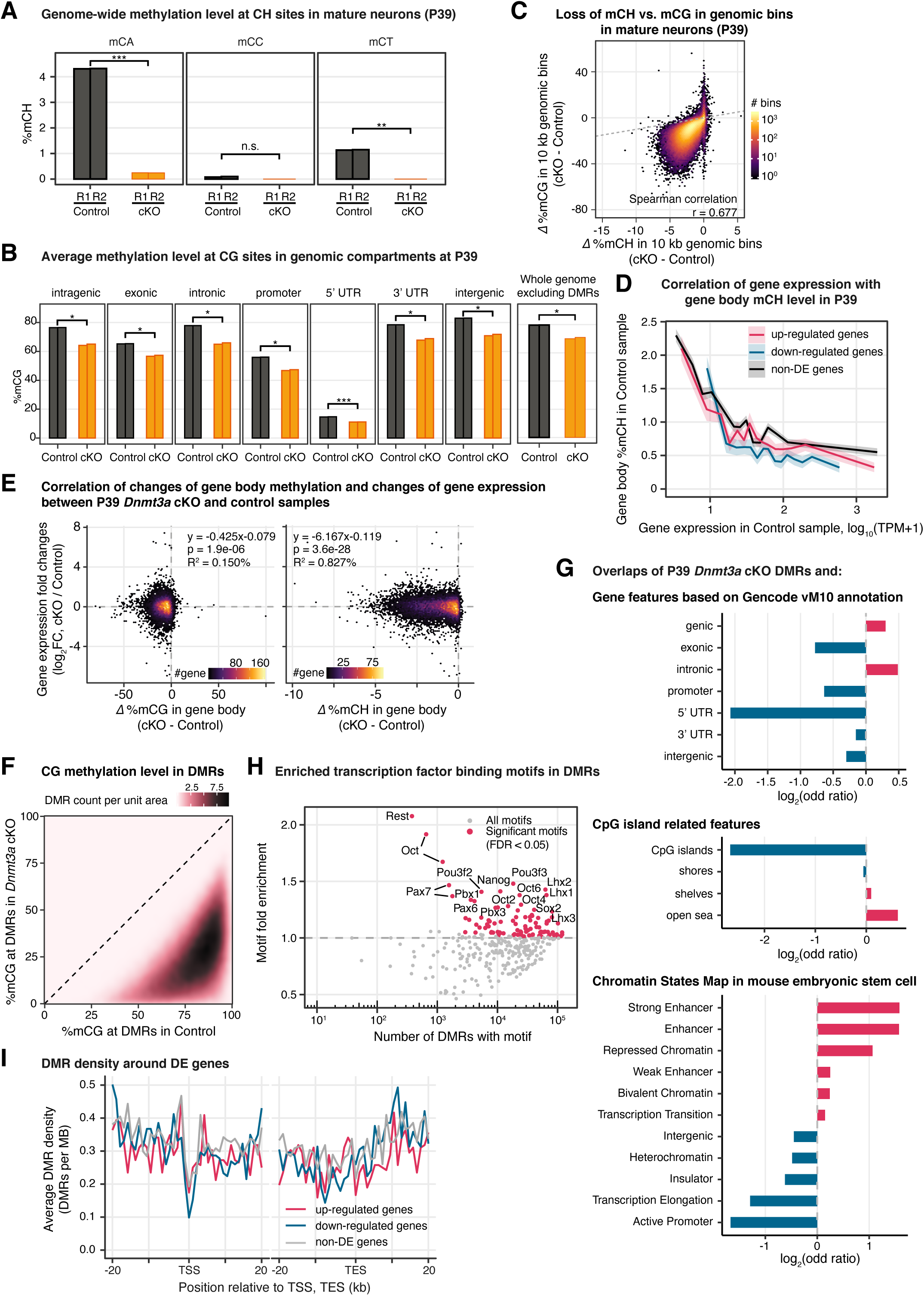
Genome-wide reduction of DNA methylation was observed in *Dnmt3a* cKO, but this cannot fully explain the disruption in the transcriptome. **(A)** The reduction of genome-wide DNA methylation level in P39 is observed in all three non-CG contexts (CA, CC and CT). t-test significance, ***, p = 0.00087; **, p = 0.0050. **(B)** DNA methylation at CG sites is reduced across all functional genomic compartments in P39. UTR, untranslated region. t-test significance, ***, p < 0.001; *, p < 0.05. **(C)** Reduced mCG is strongly correlated with the reduction in mCH in P39 in 10-kb tiling genomic bins (258,012 bins with at least 10 reads covered in each sample). r, Spearman correlation coefficient. **(D)** Correlation of gene expression and gene body mCH level for up-regulated genes (red), down-regulated genes (blue) and non-DE genes (black) in the P39 Control samples. For each gene group, genes are stratified by their expression in the control sample by 15 bins, and the mean gene body mCH levels are plotted. The shaded ribbon areas indicate the standard error of the mean. TPM, transcripts per million. **(E)** Density scatter plots showing the relationship between changes of gene body methylation (delta mCG or mCH) and the gene expression fold changes for expressed genes (14,534 genes) between P39 *Dnmt3a* cKO and control samples. The linear regression fits, p-values and variances explained by Δ%mC (R^2^) are shown. **(F)** 2-D distribution of mCG levels in P39 Control vs. *Dnmt3a* cKO samples at DMRs. DMR density is estimated through a Gaussian smoothed kernel. **(G)** Enrichment (red) or depletion (blue) of P39 cKO DMRs in GENCODE annotated gene features (top), in CpG island related features (middle), and in the chromatin states map in mouse embryonic stem cell (bottom). All enrichments and depletions shown are significant (fisher test p < 0.05). **(H)** Number of known transcription factor binding motifs within P39 *Dnmt3a* cKO hypo-DMRs and their fold enrichment. Significant motifs (FDR < 0.05) are colored in red. **(I)** The density of P39 *Dnmt3a* cKO hypo-DMRs around differentially expressed genes. TSS, Transcription Start Site; TES, Transcription End Site.

**Supplementary Figure 8.**
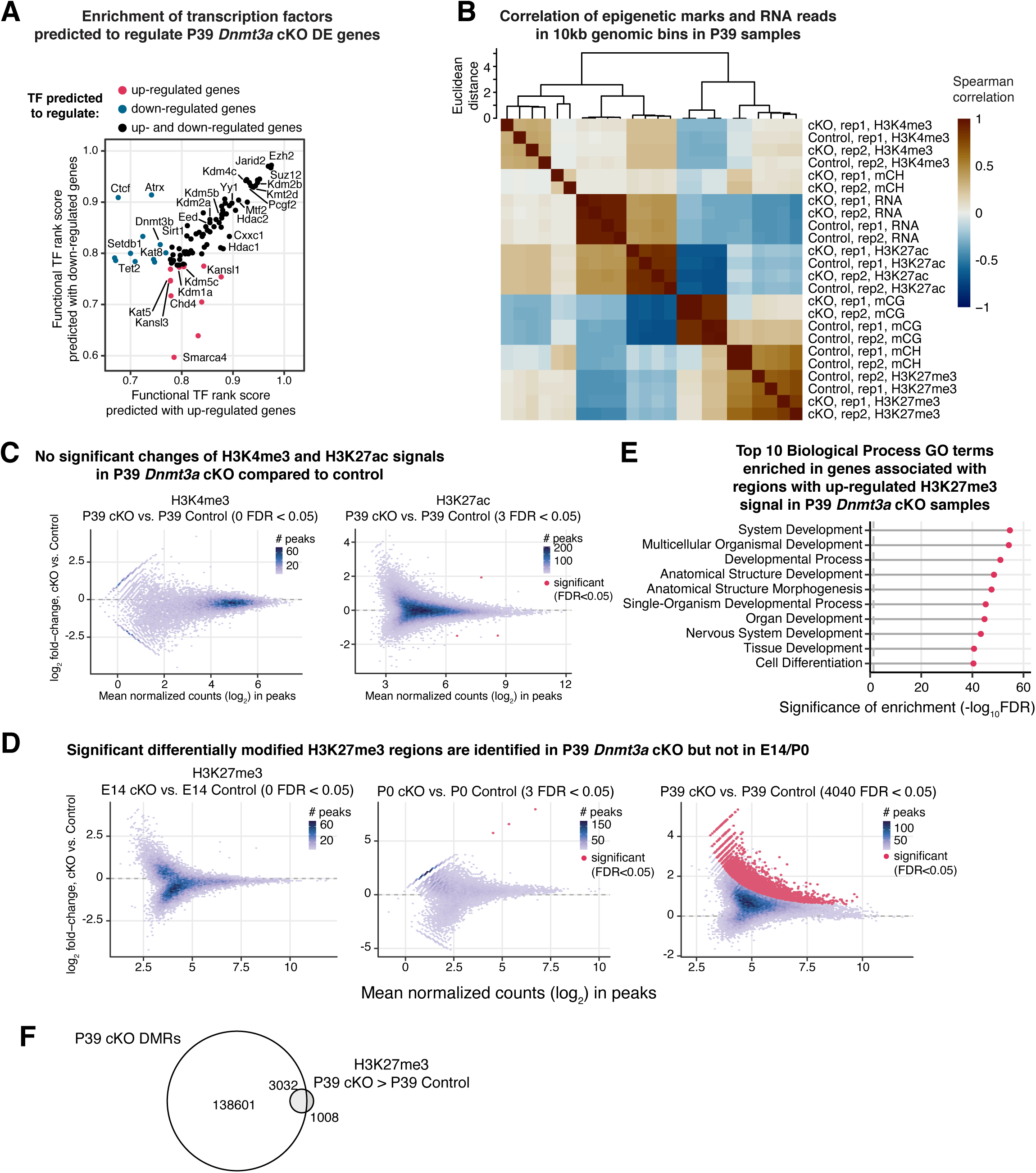
H3K27me3 signal was increased in P39 *Dnmt3a* cKO and highly overlapped with DMRs. **(A)** Transcription factors (TFs) associated with chromatin organization were predicted to regulate DE genes in the P39 *Dnmt3a* cKO. The functional TF rank score was given by Binding Analysis of Regulation of Transcription (BART)(Wang et al., 2018). TFs predicted to regulate only up-regulated genes, to regulate only down-regulated genes and to regulate both the up- and down-regulated genes are colored in red, blue and black, respectively. **(A)** Correlation heatmap of epigenetic marks and RNA reads in 10 kb tiling genomic bins. For mCG and mCH, the mean methylation in each bin is used. For RNA-seq and histone modification ChIP-seq, the RPKM (reads per kilobase per million) value in each bin is used. The dendrogram is calculated by hierarchical clustering with complete linkage using the euclidean distances of the Spearman correlation across the samples. Rep1/2, replicate 1/2. **(C)** MA plots show that no significant changes of H3K4me3 and H3K27ac signal in P39 *Dnmt3a* cKO compared to control. **(D)** MA plots show that significant differentially modified H3K27me3 regions are only found in P39 *Dnmt3a* cKO but not E14 or P0 when compared to control. **(E)** Top 10 Gene Ontology terms of the Biological Process ontology enriched in the genes associated with regions with up-regulated H3K27me3 signal in the P39 *Dnmt3a* cKO samples. Gene-region association was estimated using GREAT (McLean et al., 2010). FDR, false discovery rate; the vertical dashed line shows the threshold of FDR = 0.05. **(F)** Venn diagrams show overlaps of P39 *Dnmt3a* cKO DMRs and H3K27me3 and H3K27ac peaks in P39 Control samples. 64.6% of the H3K27me3 peaks overlap with DMRs and 22.4% of the H3K27ac peaks overlap with DMRs. **(G)** Overlaps of P39 H3K27me3 differentially modified regions with P39 *Dnmt3a* cKO DMRs.

**Supplementary Figure 9.**
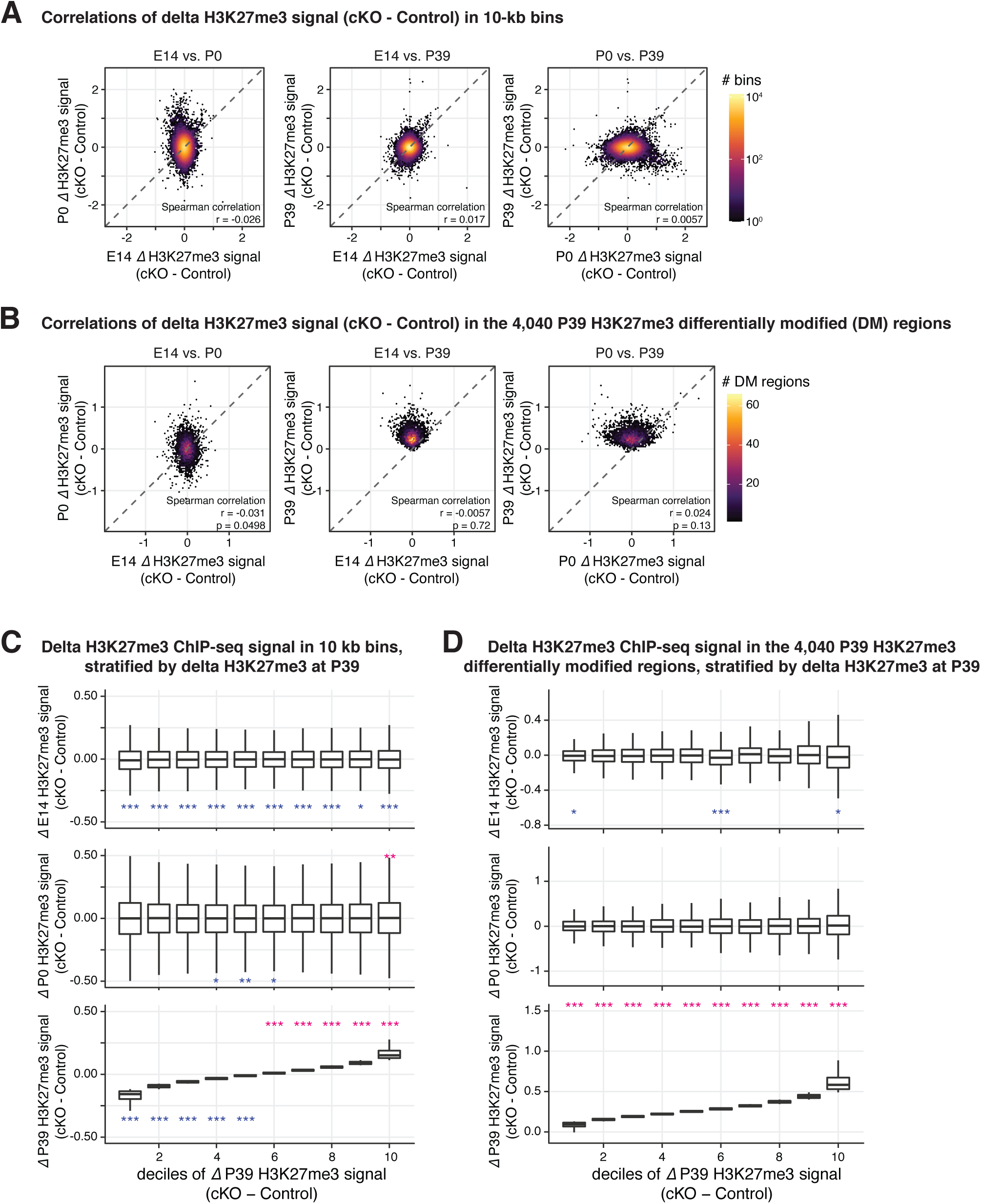
The increased H3K27me signal in P39 was generally not observed in E14 or P0. **(A-B)** Correlations of the differences of H3K27me3 signal between *Dnmt3a* cKO and control across the three development time points in 10-kb genomic bins (**A**) and the P39 H3K27me differentially modified (DM) regions (**B**). **(C-D)**, boxplots to show the distribution of the differences of H3K27me3 signal between *Dnmt3a* cKO and Control in E14, P0, and P39, stratified by ΔH3K27me3 signal at P39. Signals are counted in 10-kkb genomic bins (**C**, as in **A**) and the P39 H3K27me DM regions (**D**, as in **B**). Asterisks show the significances from Wilcoxon rank sum tests against zero (red, greater than zero; blue less than zero; *, p < 0.05; **, p < 0.01; ***, p < 0.001).

**Supplementary Figure 10.**
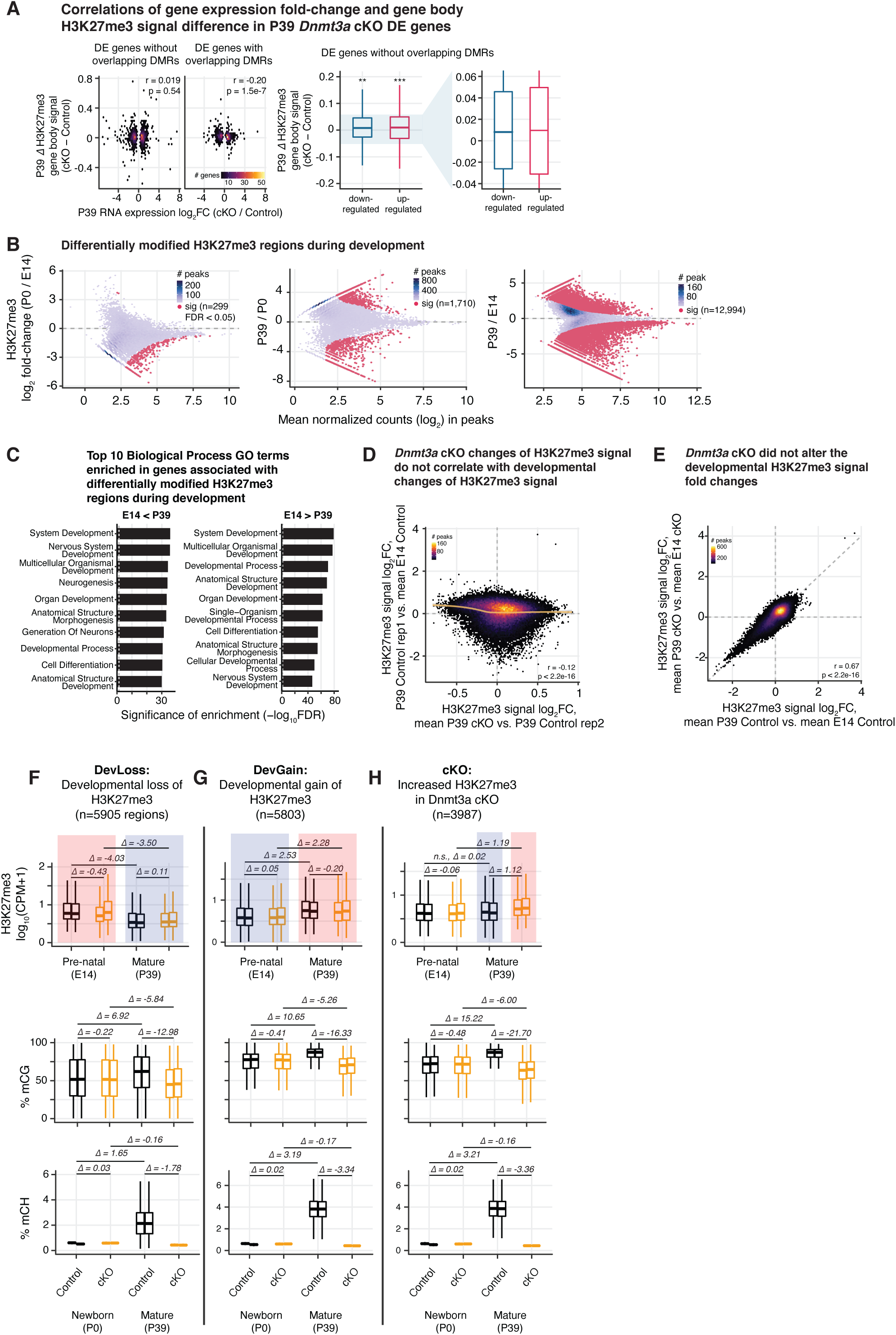
Regions prone to alteration of H3K27me3 by *Dnmt3a* cKO were distinct from the regions affected by developmentally dynamic H3K27me3. **(A)** Correlations of gene expression fold-change and gene body H3K27me3 signal difference in P39 *Dnmt3a* cKO DE genes. Left, scatter plot to show the correlation. DE genes are grouped by whether or not they overlap with P39 *Dnmt3a* cKO DMRs. r, Spearman correlation coefficient. Right, P39 *Dnmt3a* cKO DE genes without overlapping P39 cKO DMRs in general show small but significant increases of H3K27me3 in P39 cKO. Asterisks show the fold change significance against zero using Wilcoxon rank sum test (**, p < 0.01; ***, p < 0.001). Note that the effect sizes are smaller than those of DE genes with overlapping P39 cKO DMRs (related to Figure 6B). **(B)** Differentially modified H3K27me3 regions during development in control samples. **(C)** Top 10 most enriched biological process terms from the Gene Ontology in the genes associated with differentially modified H3K27me3 regions during development in control samples. Gene-region association was predicted by GREAT (McLean et al., 2010). FDR, false discovery rate; the vertical dashed line shows the threshold of FDR = 0.05. **(D)** Correlations of P39 H3K27me3 signal fold-changes of *Dnmt3a* cKO vs. Control and developmental H3K27me3 signal fold-changes of P39 vs. E14 in H3K27me3 ChIP-seq peaks. Note that peaks with big absolute fold-changes in P39 cKO vs. P39 Control are generally not changed much in P39 Control vs. E14 Control, and vice versa. Replicate 1 and 2 of P39 Control are used in the y and x axis, respectively, to avoid double dipping. The smoothed line is fitted using a generalized additive model, and the shaded area shows 95% confidence interval of the fit. r, Spearman correlation coefficient. **(E)** The developmental H3K27me3 signal fold changes are not affected by *Dnmt3a* cKO. r, Spearman correlation coefficient. **(F-H)** boxplots to show the distribution of H3K27me3 signal, mCG and mCH levels in peaks that overlaps with developmental loss-of-H3K27me3 regions **(F)**, peaks that overlaps with developmental gain-of-H3K27me3 regions **(G)** and peaks that overlaps with increase H3K27me3 regions in P39 *Dnmt3a* cKO **(H)**. Asterisks show the significances from paired t-tests (***, p < 0.001; n.s., not significant). Δ values are the mean of the differences for the two groups in comparison (group on the right vs. group on the left).

## Supplemental methods

### Generation of the *Dnmt3a* cKO mice line

All animal procedures were conducted in accordance with the guidelines of the American Association for the Accreditation of Laboratory Animal Care and were approved by the Salk Institute for Biological Studies Institutional Animal Care and Use Committee. For behavior, slice physiology and spine analyses, *Dnmt3a*-floxed animals(Okano et al., 1999) (backcrossed to C57BL/6 for at least seven generations) were crossed to *Nex*-Cre (*Neurod6*-Cre)(Goebbels et al., 2006) mice to generate *Dnmt3a*-KO animals carrying the deletion only in pyramidal cells. To be able to isolate pyramidal neuron nuclei for DNA methylation, transcription, and ChIP analyses, the mouse lines (*Dnmt3a*-KO and *Nex*-Cre) were crossed to a mouse line carrying the INTACT background (B6.129-*Gt(ROSA)26Sor^tm5(CAG-Sun1/sfGFP)Nat/MmbeJ^*). The deletion of *Dnmt3a* from pyramidal cells was confirmed by RNA-seq (deletion of exon 19) and Western blot (Figure 1B, S1B and S1C). For both backgrounds, Nex-Cre hemizygous mice were used as controls.

### Frontal cortex dissection, Nuclei isolation, and flow cytometry

Frontal cortex tissue was produced as described(Lister et al., 2013; Luo et al., 2017) from postnatal day 39 (P39) *Dnmt3a* cKO and control animals. The nuclei of GFP-expressing NeuN-positive excitatory neurons were isolated and collected using fluorescence activated nuclei sorting (FANS) as described(Lister et al., 2013; Luo et al., 2017) with the following modification: prior to FANS, nuclei were labeled with anti-NeuN-AlexaFluor647 and anti-GFP-AlexaFluor488. Nuclei were sorted as described(Lister et al., 2013). Double positive nuclei were retained for RNA-seq, ChIP-seq and MethylC-seq library preparation and sequencing.

### Western blot

Frontal cortex proteins were obtained by homogenization in RIPA buffer of the following composition: 150 mM NaCl, 10 mM Na2HPO4, 1% NaDOC, 1% NP-40, 0.5% SDS, 1 mM DTT, 1 mM PMSF in DMSO, supplemented with protease inhibitor (Sigma-Aldrich #11836153001) and phosphatase inhibitor (Pierce #A32957) cocktails. After centrifugation at 15000 x g, supernatants were preserved and protein concentration was determined by the BCA method (Pierce). Protein bands were separated in 8% PAGE-gels and transferred to nitrocellulose membranes. After blocking in TBS-tween with 5% milk, DNMT3A was detected by the use of anti-DNMT3A antibody (Abcam) and chemiluminescence. DNMT3A bands were normalized to ACTIN content in each sample.

### RNA extraction, RNA-seq library construction and sequencing

Nuclei (between 50,000-60,000) were used to isolate RNA using the Single-Cell RNA Purification Kit (Norgen, catalog# 51800). In brief, aliquots of nuclei were resuspended in 350 µl of RL buffer (Norgen) and passed through an 18G syringe five times. RNA extraction, including DNase digestion, followed manufacturers’ instructions. RNA was eluted in 20 µl of Elution Solution A (Norgen). The nuclear RNA concentration was determined using TapeStation (Agilent). RNA was diluted to 1 ng/µl and a total of 5 ng processed for RNA-seq library preparation. RNA libraries were prepared using NuGen Ovation® RNA Seq System V2 (#7102-32) for cDNA preparation following the product manual. cDNA purification was done using Zymo Research DNA Clean & Concentrator™-25 with modification from Ovation protocol. cDNA, eluted in 30-40 µl of TE (1 µg per sample), was fragmented at 300bp using Covaris S2 (Sonolab S-series V2), followed by library preparation according to KAPA LTP Library Preparation Kit (KK8232), using Illumina indexed adapters.

### DNA extraction

DNA extraction was performed using the Qiagen DNeasy Blood and Tissue kit (catalog #69504) and eluted into 50-100 µL AE.

### Genomic DNA library construction and sequencing

1.5 µg of genomic DNA was fragmented with a Covaris S2 (Covaris, Woburn, MA) to 400 bp, followed by end repair and addition of a 3’ A base. Cytosine-methylated adapters provided by Illumina (Illumina, San Diego, CA) were ligated to the sonicated DNA at 16°C for 16 hours with T4 DNA ligase (New England Biolabs). Adapter-ligated DNA was isolated by two rounds of purification with AMPure XP beads (Beckman Coulter Genomics, Danvers, MA). Half of adapter-ligated DNA molecules were enriched by 6 cycles of PCR with the following reaction composition: 25 µL of Kapa HiFi Hotstart Readymix (Kapa Biosystems, Woburn, MA) and 5 µl TruSeq PCR Primer Mix (Illumina) (50 µl final). The thermocycling parameters were: 95°C 2 min, 98°C 30 sec, then 6 cycles of 98°C 15 sec, 60°C 30 sec and 72°C 4 min, ending with one 72°C 10 min step. The reaction products were purified using AMPure XP beads and size selection done from 400 - 600 bp. Libraries were sequenced on Illumina HiSeq 4000.

### MethylC-seq library construction and sequencing

MethylC-seq libraries were prepared as previously described(Urich et al., 2015). All DNA obtained from the extraction was spiked with 0.5% unmethylated Lambda DNA. The DNA was fragmented with a Covaris S2 (Covaris, Woburn, MA) to 200 bp, followed by end repair and addition of a 3’ A base. Cytosine-methylated adapters provided by Illumina (Illumina, San Diego, CA) were ligated to the sonicated DNA at 16°C for 16 hours with T4 DNA ligase (New England Biolabs). Adapter-ligated DNA was isolated by two rounds of purification with AMPure XP beads (Beckman Coulter Genomics, Danvers, MA). Adapter-ligated DNA (:S450 ng) was subjected to sodium bisulfite conversion using the MethylCode kit (Life Technologies, Carlsbad, CA) as per manufacturer’s instructions. The bisulfite-converted, adapter-ligated DNA molecules were enriched by 8 cycles of PCR with the following reaction composition: 25 µL of Kapa HiFi Hotstart Uracil+ Readymix (Kapa Biosystems, Woburn, MA) and 5 µl TruSeq PCR Primer Mix (Illumina) (50 µl final). The thermocycling parameters were: 95°C 2 min, 98°C 30 sec, then 8 cycles of 98°C 15 sec, 60°C 30 sec and 72°C 4 min, ending with one 72°C 10 min step. The reaction products were purified using AMPure XP beads. Up to two separate PCR reactions were performed on subsets of the adapter-ligated, bisulfite-converted DNA, yielding up to two independent libraries from the same biological sample. MethylC-seq libraries were sequenced on Illumina HiSeq 4000.

### ChIP-seq library construction and sequencing

Sorted nuclei were crosslinked for 15min in 1% formaldehyde solution and quenched afterward with glycine at a final concentration of 0.125 M. After crosslinking, nuclei were sonicated in Lysis buffer (50 mM Tris HCl pH 8, 20 mM EDTA, 1% SDS, 1X EDTA free protease inhibitor cocktail). ChIP assays were conducted with antibodies against H3K27me3 (39156, Active Motif), H3K27ac (39133, Active Motif) and H3K4me3 (04-745, Millipore Sigma). Mouse IgG (015-000-003, Jackson ImmunoResearch) served as negative control. H3K4me3 ChIP-seq assays were conducted with 100K nuclei and 500K nuclei were used for H3K27me3 and H3K27ac ChIP-seq assays. The respective antibodies and IgG were coupled for 4-6 hours to Protein G Dynabeads (50 µl, 10004D, Thermo Fisher Scientific). Equal amounts of sonicated chromatin were diluted with 9 volumes of Binding buffer (1% Triton X-100, 0.1% Sodium Deoxycholate, 2X EDTA free protease inhibitor cocktail) and subsequently incubated overnight with the respective antibody-coupled Protein G beads. Beads were washed successively with low salt buffer (50 mM Tris HCl pH 7.4, 150 mM NaCl, 2 mM EDTA, 0.5% Triton X-100), high salt buffer (50 mM Tris HCl pH 7.4, 500 mM NaCl, 2 mM EDTA, 0.5% Triton X-100) and wash buffer (50 mM Tris HCl pH 7.4, 50 mM NaCl, 2 mM EDTA) before de-crosslinking, proteinase K digestion and DNA precipitation. Libraries were generated with the Accel-NGS 2S Plus DNA Library Kit (21024, Swift Biosciences) and sequenced on the Illumina HiSeq 4000 Sequencing system.

### RNA-seq data processing

RNA-seq reads were first went through quality control using FastQC(Andrews et al., 2012) (v0.11.8, https://www.bioinformatics.babraham.ac.uk/projects/fastqc/), and then were trimmed to remove sequencing adapters and low-quality sequences (minimum Phred score 20) using Trim Galore (v0.5.0, https://www.bioinformatics.babraham.ac.uk/projects/trim_galore/, a wrapper tool powered by Cutadapt(Martin, 2011) v1.16) in the paired-end mode. Clean reads were then mapped to the mouse mm10 (GRCm38) genome and the GENCODE annotated transcriptome (release M10) with STAR(Dobin et al., 2013) (Spliced Transcripts Alignment to a Reference, v2.5.1b). Gene expression was estimated using RSEM(Li and Dewey, 2011) (RNA-Seq by Expectation Maximization, v1.2.30). Gene-level “expected count” from the RSEM results were rounded and fed into edgeR(Robinson et al., 2010) (v3.24.1) to call differentially expressed genes. Only genes that were expressed (with counts per million > 2) in at least two samples were kept. These counts were then normalized using the TMM method(Robinson and Oshlack, 2010), and DE genes were then called in the quasi-likelihood F-test mode, requiring FDR < 0.05 and FC > 20% (Table S2).

### Enrichment test of Gene Ontology (GO) terms in differentially expressed genes

GO enrichment analysis was performed using clusterProfiler(Yu et al., 2012) (v3.10.0). Only “Biological Process” terms with no less than 10 genes and no more than 250 genes were considered. Terms with FDR < 0.05 were considered significantly enriched.

### Genomic DNA sequencing data processing and SNP calling

To estimate the completeness of the inbreeding of the mouse strains, and to avoid incorrect cytosine context assignment in the following methylC-seq data processing, we used the genomic DNA sequencing data of both the *Dnmt3a* cKO and the control animals to call SNPs against the mouse mm10 genome. We followed the GATK “best practices for germline SNPs and indels in whole genomes and exomes” pipeline(DePristo et al., 2011; McKenna et al., 2010; Van der Auwera et al., 2013). Briefly, raw data were first trimmed to remove sequencing adapters and low-quality sequences (minimum Phred score 20) using Trim Galore in the paired-end mode. Clean data were then mapped to the mm10 genome using BWA(Li and Durbin, 2009) (v0.7.13-r1126). Duplicates reads were marked with Picard(Broad Institute, 2018). Then the analysis-ready reads were fed into GATK (v3.7) to perform two rounds of joint genotyping and base recalibration. Variants were then filtered using the following criteria: QD < 2.0 II FS > 60.0 II MQ < 40.0 II MQRankSum < −12.5 II ReadPosRankSum < −8.0 II SOR > 4.0. By that, we identified 548,530 and 507,669 SNPs (relative to mm10) in the *Dnmt3a* cKO and the control animals, respectively. At last, we created a substituted genome to mask out all these SNPs (replaced with Ns) with the maskfasta tool in the BEDTools suite(Quinlan and Hall, 2010) (v2.27.1). This substituted genome was used in the following methylC-seq data processing pipeline.

### MethylC-seq data processing

MethylC-seq reads were processed using the methylpy pipeline (v1.3.2, https://github.com/yupenghe/methylpy) as previously described(Lister et al., 2013; Mo et al., 2015). Briefly, computationally bisulfite-converted genome index was built using the aforementioned substituted genome file appended with the lambda phage genomic sequence. MethylC-seq raw reads were first trimmed to remove sequencing adapters and low-quality sequences (minimum Phred score 10) using Cutadapt in paired-end mode. To acquire higher mappability, we treated the two ends of the clean reads as they were sequenced in single-end mode, and mapped them to the converted genome index with bowtie2(Langmead and Salzberg, 2012) (v2.3.0) as aligner in the single-end pipeline of methylpy. Only reads uniquely mapped were kept, and clonal reads were removed. The bisulfite non-conversion rate was estimated using the spiked-in unmethylated lambda phage DNA. For each cytosine, a binomial test was performed to test whether the methylation levels are significantly greater than 0 with an FDR threshold of 0.01.

For a particular genomic region, the raw methylation level for a given cytosine context (CG or CH) was defined as:

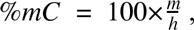

where m is the total number of methylated based calls within the region, and h is the total number of covered based calls within the region. Methylation levels were then corrected for non-conversion rate (NCR) using the following maximum likelihood formula:

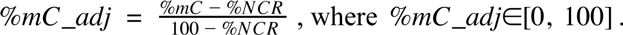

When profiling the methylation landscapes around differentially expressed genes, we selected control genes with comparable gene expression using the R package MatchIt(Ho et al., 2011) (v3.0.2). These control genes were defined with nearest neighbor matching of the expression using logistic link propensity score as a distance measure, requiring the standard deviation of the distance to be less than 0.01.

### Differentially methylated regions (DMRs) calling

CG DMRs were identified using a previously reported method(Ma et al., 2014; Schmitz et al., 2013; Schultz et al., 2015), which is implemented in the DMRfind function in methylpy. We required at least 3 differentially methylated sites within a particular DMR, and the maximum distance two significant sites can be to be included in the same DMR to be 500 nucleotides. With an FDR cutoff of 0.01 and a post-filtering cutoff of methylation levels change greater than 30%, we found 141,633 *Dnmt3a* cKO hypo-DMRs in P39 (Table S4). Note that with these criteria we also found 19 *Dnmt3a* cKO hyper-DMRs, which we thought were noise and/or SNPs failed to be detected by the masking pipeline described earlier. Therefore we removed these hyper-DMRs from further consideration.

### Enrichment test of DMRs and other genomic regions

To test whether DMRs were significantly enriched in certain genomic features, we use methods adapted from a recent report(Rizzardi et al., 2019). Briefly, for each genomic feature, we constructed a 2 by 2 contingency table of (n_11_, n_12_, n_21_, n_22_), where:

- n_11_ is the number of CG sites in DMRs that were inside the feature;
- n_12_ is the number of CG sites in DMRs that were outside of the feature;
- n_21_ is the number of CG sites not in DMRs that were inside in feature;
- n_22_ is the number of CG sites not in DMRs that were outside of the feature.

The total number of CG sites in consideration was the number of autosomal and chromosome X CG in the reference genome. Counting number of CG rather than the number of DMRs or bases accounts for the non-uniform distribution of CG along the genome and avoids double-counting DMRs that are both inside and outside of the feature.

With this contingency table, we estimated the enrichment log odd ratio (OR) along with its standard error (se) and 95% confidence interval (ci) with the following formulas:

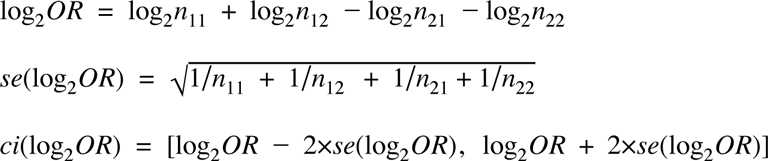

P-value from performing Fisher’s exact test for testing the null of independence of rows and columns in the contingency table (the null of no enrichment or depletion) was computed using the fisher.test() function in R.

The genomic regions/features used in these enrichment tests includes: a list of developmental DMRs that gain or lose methylation during development(Lister et al., 2013); gene features (genic, exonic, intronic, promoter, 5’UTR, 3’UTR, intergenic) based on the Gencode vM10 annotation (promoters were defined as the 2kb regions around transcription start sites); CpG island (CGI) related features based on the “cpgIslandExt” annotation from the UCSC genome browser(Karolchik et al., 2004; Kent et al., 2002) (http://genome.ucsc.edu/index.html), where CGI shores were defined as CGI 2kb, CGI shelves were defined as 2-4kb of CGI and open seas were defined as regions that were at least 4kb away from any CGI; the 12 states of the chromatin states map in mouse embryonic stem cell(Pintacuda et al., 2017) (https://github.com/guifengwei/ChromHMM_mESC_mm10) generated by ChromHMM(Ernst and Kellis, 2012, 2017) using ChIP-seq data from the ENCODE project(Davis et al., 2018; ENCODE Project Consortium, 2012); the H3K4me3, H3K27ac and H3K27me3 peaks and the H3K27me3 differentially binding regions generated with our ChIP-seq data (see the “ChIP-seq data processing” section).

### Predicting functional transcription factors regulating the differentially expressed genes with BART

The lists of up-regulated and down-regulated genes were fed into BART(Wang et al., 2016, 2018) (Binding Analysis for Regulation of Transcription) separately to predict functional transcription factors (TFs) and chromatin regulators that bind at cis-regulatory regions of the DE genes. To make a better visualization we transformed the relative rank (a metric generated by BART to represent the average rank of Wilcoxon P-value, Z-score and max AUC for each factor divided by the total number of factors) into the functional TF rank score (which is simply 1 minus the relative rank) so that the higher the rank score the more possible the TF regulates the DE genes. The integrative rank significance was estimated with the Irwin-Hall p-value. TFs with Irwin-Hall p-value < 0.05 were considered significant.

### Enrichment test of known transcription factor binding motifs in DMRs

We used the “findMotifsGenome.pl” tool in HOMER (Heinz et al., 2010) (Hypergeometric Optimization of Motif EnRichment, v4.8.3) to find known transcription factor binding motif in the *Dnmt3a* cKO DMRs. The parameters used are as follows: “-size 500 -len 8,10,12 -S 25 -fdr 100 -p 10 -mset vertebrates -bits -gc -nlen 3 -nomotif”. A set of non-neural DMRs(Hon et al., 2013) was used as the background.

### ChIP-seq data processing

ChIP-seq reads were pre-processed with the ENCODE Transcription Factor and Histone ChIP-Seq processing pipeline (https://github.com/ENCODE-DCC/chip-seq-pipeline2, v1.1.6). Briefly, paired-end reads were mapped to the mm10 genome with BWA(Li and Durbin, 2009) (v0.7.13-r1126). Reads were then filtered using samtools(Li et al., 2009) (v1.2) to remove unmapped, mate unmapped, not primary alignment and duplicates reads (-F 1804). Properly paired reads were retained (-f 2). Multi-mapped reads (MAPQ < 30) were removed. PCR duplicates were removed using the MarkDuplicates tool in Picard(Broad Institute, 2018) (v2.10.6). Reads mapped to the blacklist regions(ENCODE Project Consortium, 2012) in mouse mm10 genome (http://mitra.stanford.edu/kundaje/akundaje/release/blacklists/mm10-mouse/mm10.blacklist.bed.gz) were also removed.

Peak calling was performed using epic2(Stovner and Sretrom, 2019) (v0.0.16), a reimplementation of SICER(Zang et al., 2009). For H3K4me3 and H3K27ac, we used the following parameters: “--bin-size 200 --gaps-allowed 1”. For H3K27me3, we used the following parameters: “--bin-size 200 --gaps-allowed 3”. The IgG sample was used as a control.

Differentially modified regions of the histone modification ChIP-seq data were called using DiffBind(Ross-Innes et al., 2012) (v2.10.0) in DESeq2(Love et al., 2014) (v1.22.1) mode. Regions with FDR < 0.05 were considered significant. Genes associated with these regions were identified using GREAT(McLean et al., 2010) (v3.0.0) with the “Basal plus extension” association rule with default parameters. GO enrichment analysis of the associated genes was performed with GREAT, and we considered the GO biological process terms with hypergeometric test FDR < 0.05 as significant.

### Definition of bivalent and active CGI promoters

CGI promoters were defined as CpG islands (downloaded from the UCSC genome browser) that overlapping with promoters (±2kb regions around transcription start sites annotated in Gencode vM10). These CGI promoters were further tested to see whether they overlapped with the ChIP-seq peaks of H3K4me3 and H3K27me3. Bivalent CGI-promoters were defined as CGI-promoters that overlapped with both the H3K4me3 and H3K27me3 peaks, whereas active CGI-promoters were defined as CGI-promoters that overlapped with only the H3K4me3 peaks but not the H3K27me3 peaks.

### Identification of DNA methylation valleys (DMVs)

To find DMVs, we first identified UMRs (undermethylated regions) and LMRs (low methylated regions) using MethylSeekR(Burger et al., 2013) (v1.22.0) with m = 0.5, n = 7 and FDR < 0.05. Noted that MethylSeekR did not recover any PMDs (partially methylated domains). DMVs were then defined as UMRs that with length ≥ 5 kb and mean methylation level : ≤ 15%. To compare DMVs identified in the *Dnmt3a* cKO and the control samples, we further grouped these DMVs into 5 categories, namely consistent (exact same DMV in the two conditions), cKO unique, control unique, expanded (wider in the cKO) and shrunken (wider in the control).

### DMR enrichment around CGI promoter

We used regioneR(Gel et al., 2016) (v1.14.0) to test whether two sets of genomic regions had significantly higher numbers of overlaps compared to expected by chance. We used permTest() to perform the permutation test, and used the randomizeRegions() function to generate the shuffled control for 5,000 times, where the query regions were randomly placed along the genome independently while maintaining their size. The strength of the association of the two sets of regions was estimated using z-score, the distance (measured in standard deviation) between the expected overlaps in the shuffled control and the observed overlaps, and the p-value was reported. To check if the association was specifically linked to the exact position of the query regions, we used the localZscore() function with window = 5000 and step = 50, which shifted the query regions and estimated how the value of the z-score changed when moving the regions.

### Other tools used in the data analysis

Browser representations were created using AnnoJ(Lister et al., 2009). All analyses were conducted in R (v3.5.0) or Matlab 2017a. Genomic ranges manipulation was done either with bedtools(Quinlan and Hall, 2010) or GenomicRanges(Lawrence et al., 2013). Multiple comparison correction for p-values was performed with the Benjamini-Hochberg FDR method(Benjamini and Hochberg, 1995). Results with FDR < 0.05 were considered significant except where stated otherwise. The smoothed lines in Figure 6C and S10C were fitted with a generalized additive model using the “gam” function in the “mgcv” R package, with formula = y ∼ s(x, bs = “cs”).

### Patch-clamp electrophysiology

Male and female mice (6-9 weeks) were anesthetized with isoflurane and decapitated. The brains were quickly removed and coronal slices of the frontal cortex containing the prelimbic region (∼2 mm anterior to bregma) were cut in ice-cold slicing medium of the following composition (in mM): 110 sucrose, 2.5 KCl, 0.5 CaCl_2_, 7 MgCl_2_, 25 NaHCO_3_, 1.25 NaH_2_PO_4_, and 10 glucose (bubbled with 95% O_2_ and 5% CO_2_). The slices were then transferred to artificial CSF (aCSF) containing (in mM) 130 NaCl, 2.5 KCl, 1.25 NaH_2_PO_4_, 23 NaHCO_3_, 1.3 MgCl_2_, 2 CaCl_2_ and 10 glucose, equilibrated with 95% O_2_ and 5% CO_2_ at 35°C for 30 minutes and afterward maintained at room temperature (22-24 °C) for at least 1 hour (patch-clamp recording) before use. Brain slices were then transferred to a recording chamber and kept minimally submerged under continuous superfusion with aCSF at a flow rate of ∼2 ml/min. Whole-cell recordings were obtained from putative prelimbic layer 2 (L2) pyramidal cells (identified by their pyramidal-shaped cell bodies and long apical dendrite using an upright microscope equipped with differential interference contrast optics). In acute mPFC slices, the prelimbic L2 is clearly distinguishable from L1 and L3 as a thin dark band that is densely packed with neuron somata. Pipettes had a tip resistance of 4-8 MO when filled with an internal solution of the following composition (in mM): 125 K-gluconate, 15 KCl, 8 NaCl, 10 HEPES, 2 EGTA, 10 Na_2_Phosphocreatine, 4 MgATP, 0.3 NaGTP (pH 7.25 adjusted with KOH, 290-300 mOsm). Access resistance (typically 15-35 Mn) was monitored throughout the experiment to ensure stable recordings.

After obtaining the whole-cell configuration in voltage clamp mode, cells were switched from a holding potential of −70 mV to current-clamp mode and the bridge-balance adjustment was performed. Passive electrical properties were quantified from recordings with hyperpolarizing current injections that evoked small −5 mV deflections in membrane potential from resting. Responses to stepwise current injections (10-300 pA in increments of 10 pA; duration, 1 s) were recorded at 20 kHz in order to calculate input-output curves and rheobase - the minimal current necessary to trigger the first action potential. Miniature excitatory postsynaptic currents (mEPSCs) were recorded for 5 minutes in voltage-clamp mode (Vh=-70 mV) in the presence of the Na+ channel blocker, TTX (0.5 µM), to prevent the generation of action potentials, and picrotoxin (50 µM), an antagonist of GABA_A_ receptors, to minimize inhibitory responses. In these conditions, mEPSCs could be blocked by the AMPA receptor antagonist, CNQX (25 µM). Single events larger than 6 pA were detected off-line using the Minianalysis program (Synaptosoft Inc. Decatur, GA). All data were acquired using a Multiclamp 700B amplifier and pCLAMP 9 software (Molecular Devices, LLC. San Jose, CA).

### Fluorescent labeling of dendritic spines

Coronal brain slices containing the mPFC of 10-13 weeks female mice were prepared as for electrophysiological recordings and placed in a beaker for 3 hours at room temperature (24° C) to allow functional and morphological recovery. One slice was then transferred to a recording chamber and kept minimally submerged under continuous superfusion with aCSF bubbled with carbogen (95% O_2_ / 5% CO_2_) at a flow rate of −2 ml/min. Previously sonicated crystals of the fluorescent marker DiI were placed next to the somata of layer 2 neurons in the prelimbic cortex, identified with the aid of an upright microscope equipped with differential interference contrast optics. The mice in these experiments had a Thy1-YFP background to help rule out non specific labeling of deeper layer neurons. The neurons were exposed to the DiI crystals for 60 minutes. The slices were then gently removed from the incubation chamber with a transfer pipette and immersed in fixative (4% PFA) for 30 minutes. Then, the slices were rinsed three times with PBS for 5-10 minutes each, after which they were mounted on slides with prolong gold antifade mounting medium (Life Technologies - Molecular Probes). The slides were kept in a dark box for 24 h at room temperature to allow the liquid medium to form a semi-rigid gel. Imaging took place 24-48 h from the time of the initial staining.

### Confocal imaging

Dendritic spines were imaged by an investigator blind to the genotype using a Zeiss AiryScan confocal laser scanning microscope. All images were taken using the Zeiss Plan-APOCHROMAT 63x oil-immersion lens (N/A 1.4). A 543nm laser was used to visualize the fluorescence emitted by DiI. Serial stack images with a 0.2 µm step size were collected, and then projected to reconstruct a three dimensional image that was post-processed by the AiryScan software. Dendritic segments in layer 1, which were derived from layer 2 pyramidal neurons retrogradely labeled with DiI and that were well separated from neighboring neural processes, were randomly sampled and imaged. Each dendritic segment imaged for quantification belonged to a different neuron.

### Dendritic spine quantification

The z-stack series were imported into the Reconstruct software (http://synapseweb.clm.utexas.edu/software-0/), with which a second investigator also blind to the genotype performed the identification of dendritic spines and their morphometric analysis. By scrolling through the stack of different optical sections, individual spine heads could be identified with greater certainty. All dendritic protrusions with a clearly recognizable stalk were counted as spines. Spine density was determined by summing the total number of spines per dendritic segment length (30-40 µm) and then calculating the average number of spines per µm. Individual dendritic spines were classified in the following order according to pre-established criteria: protrusion longer than 3 µm, filopodia; head wider than 0.6 µm, mushroom; protrusion longer than 2 µm and head narrower than 0.6 µm, long-thin; protrusion longer than 1 µm and head narrower than 0.6 µm, long-thin; the remaining spines were labeled stuby). Branched spines (with more than one neck) were counted separately.

### BEHAVIORAL TESTING

Phenotypic characterization was initiated when the animals reached 9 weeks of age using cohorts of 10-15 male or female mice per genotype, according to the order described below.

#### Open field test

The open field test was performed using MED Associates hardware and the Activity Monitor software according to the manufacturer’s instructions (MED Associates Inc, St. Albans, VT). Animals were individually placed into clear Plexiglas boxes (43.38 x 43.38 x 30.28 cm) surrounded by multiple bands of photo beams and optical sensors that measure horizontal and vertical activity. Movement was detected as breaks within the beam matrices and automatically recorded for 60 minutes.

#### Light/dark transfer test

The light/dark transfer procedure was used to assess anxiety-like behavior in mice by capitalizing on the conflict between exploration of a novel environment and the avoidance of a brightly lit open field (150-200 lux in our experiments). The apparatus were Plexiglas boxes as for the open field test (43.38 x 43.38 x 30.28 cm) containing dark box inserts (43.38 x 12.8 x 30.28 cm). The compartments were connected by an opening (5.00 x 5.00 cm^2^) located at floor level in the center of the partition. The time spent in the light compartment was used as a predictor of anxiety-like behavior, i.e. a greater amount of time in the light compartment was indicative of decreased anxiety-like behavior. Mice were placed in the dark compartment (4-7 lux) at the beginning of the 15 minute test.

#### Elevated plus maze

The maze consisted of four arms (two open without walls and two with enclosed walls) 30 cm long and 5 cm wide in the shape of a plus sign. The apparatus was elevated approximately 33 cm over a table. At the beginning of each trial, one animal was placed inside a cylinder located at the center of the maze for 1 min. The mouse was then allowed to explore the maze for 5 minutes. The session was video-recorded by an overhead camera and subjected to automated analysis using ANY-maze software. The apparatus was wiped down with sani-wipes between trials to remove traces of odor cues. The percentage of time spent in open or closed arms was scored and used for analysis.

#### Y maze test for spontaneous alternations

Spontaneous alternations between three 38 cm long arms of a Y-maze were taken as a measure of working memory. Single 6 minute trials were initiated by placing each mouse in the center of the Y maze. Arm entries were recorded with a video camera and the total number of arm entries, as well as the order of entries, was determined. The apparatus was wiped down with sani-wipes between trials to remove traces of odor cues. Spontaneous alternations were defined as consecutive triplets of different arm choices and % spontaneous alternation defined as number of spontaneous alternations divided by the total number of arm entries minus 2.

#### Social Approach

The apparatus consisted of a Plexiglas box (60 x 38 x 23.5 cm) divided into 3 compartments by Plexiglas partitions containing openings through which the mice could freely enter the 3 chambers. The test was conducted in two 10-minute phases. In phase I, the test mouse is first allowed to explore the chambers for 10 minutes. Each of the two outer chambers contained an empty, inverted stainless steel wire cup. In phase II, the test mouse is briefly removed, and a sex-matched unfamiliar mouse, was placed under one of the wire cups and plastic blocks were placed under the other wire cup. The test mouse was then gently placed back in the arena and given an additional 10 minutes to explore. An overhead camera and video tracking software (ANY-maze, Wood Dale, IL) were used to record the amount of time spent in each chamber. The location (left or right) of the novel object and novel mouse alternates across subjects.

#### Acoustic startle responses and prepulse inhibition of the acoustic startle response

Acoustic startle responses were tested inside SR-LAB startle apparatus (San Diego Instruments, San Diego, CA), consisting of an inner chamber with a speaker mounted to the wall and a cylinder mounted on a piezoelectric sensing platform on the floor. At the beginning of testing, mice were placed inside the cylinder and then were subjected to background 65 dB white noise during a 5-min acclimation period. The PPI session began with the presentation of six pulse-alone trials of 120 dB, 40 ms. Then, a series of pulse-alone trials and prepulse trials (69, 73, or 81 dB, 20 ms followed by 100 ms pulse trial, 120 dB) were each presented 12 times in a pseudorandom order. The session concluded with the presentation of six pulse-alone trials. The apparatus was wiped down with sani-wipes between trials to remove traces of odor cues. The startle amplitude was calculated using arbitrary units, and the acoustic startle response was the average startle amplitude of pulse-alone trials. The percent PPI was calculated as follows: [100- (mean prepulse response/mean pulse alone response) x 100].

#### Cued and contextual fear conditioning

Fear conditioning experiments were performed using automated fear conditioning chambers (San Diego Instruments, San Diego, CA), similarly to previous studies(Gresack et al., 2010; Risbrough et al., 2014). On day 1, after a 2-min acclimation period, mice were presented with a tone conditioned stimulus (CS: 75 dB, 4 kHz) for 20 s that co-terminated with a foot shock unconditioned stimulus (1 s, 0.5 mA). A total of three tone-shock pairings were presented with an inter-trial interval of 40 s. To assess acquisition, freezing was quantified during foot shock presentations. Mice were returned to their home cages 2 min after the final shock. These moderate shock parameters were previously found suitable to detect both increases and decreases in fear-conditioned behavior(Risbrough et al., 2014). 24 h later, on day 2, mice were re-exposed to the conditioning chamber to assess context-dependent fear retention. This test lasted 8 min during which time no shocks or tones were presented and freezing was scored for the duration of the session. Time freezing was quantified across four 2-min blocks. Day 3: 24 h after the context fear-retention test, mice were tested for CS-induced fear retention and extinction. The context of the chambers was altered across several dimensions (tactile, odor, visual) for this test in order to minimize generalization from the conditioning context. After a 2 min acclimation period, during which time no tones were presented (’pre-tone’), 32 tones were presented for 20 s with an inter-trial interval of 5 s. Freezing was scored during each tone presentation and quantifications were done in eight blocks of four tones. Mice were returned to their home cage immediately after termination of the last tone. On day 4, after a 2 min acclimation period, during which time no tones were presented (’pre-tone’), a shorter session of 16 tones was used to assess extinction. Time freezing was quantified across four blocks of four tones.

## Supplementary Tables

**Supplementary Table 1. Sequencing metrics for RNA-seq, MethylC-seq, ChIP-seq and genomic DNA for the Dnmt3a cKO and control samples.**

**Supplementary Table 2. Gene expression and list of differentially expressed genes in P39 Dnmt3a cKO and P39 Control.**

**Supplementary Table 3. List of enriched Gene Ontology terms for biological process in differentially expressed genes (P39 Dnmt3a cKO vs. control)**

**Supplementary Table 4. List of differentially methylated regions (DMRs).**

**Supplementary Table 5. Known transcription factor motif enrichment in P39 Dnmt3a cKO DMRs**

**Supplementary Table 6. List of transcription factors and chromatin regulators that bind at cis-regulatory regions of the differentially expressed genes in P39 Dnmt3a cKO**

**Supplementary Table 7. List of ChIP-seq peaks.**

**Supplementary Table 8. List of up-regulated H3K27me3 signal regions in P39 Dnmt3a cKO.**

**Supplementary Table 9. List of enriched Gene Ontology terms for biological process in genes associated with up-regulated H3K27me3 regions in P39 Dnmt3a cKO.**

**Supplementary Table 10. List of developmental regulated H3K27me3 signal regions in control pyramidal neurons.**

**Supplementary Table 11. List of bivalent and active CGI promoters.**

**Supplementary Table 12. List of DNA methylation valleys (DMVs).**

## Notes

### Competing Interest Statement

The authors have declared no competing interest.

### Summary of Updates

New data from chromatin immunoprecipitation-sequencing (ChIP-Seq) experiments performed at developmental time points (embryonic day 14 and postnatal day 0) were added. The results were revised to incorporate these new data and to describe the developmental trajectory of changes in H3K27me3 in Dnmt3a cKO.

